# Enhancement of Target Specificity of CRISPR-Cas12a by Using a Chimeric DNA-RNA Guide

**DOI:** 10.1101/2020.02.04.933614

**Authors:** Hanseop Kim, Wi-jae Lee, Seung-Hun Kang, Junho K. Hur, Hyomin Lee, WooJeung Song, Kyung-Seob Lim, Young-Ho Park, Bong-Seok Song, Yeung Bae Jin, Bong-Hyun Jun, Dong-Seok Lee, Sun-Uk Kim, Seung Hwan Lee

## Abstract

The CRISPR-Cas9 system is widely used for target-specific genome engineering. Cpf1 is one of the CRISPR effectors that controls target genes by recognizing thymine-rich protospacer adjacent motif (PAM) sequences. Cpf1 has a higher sensitivity to mismatches in the guide RNA than does Cas9; therefore, off-target sequence recognition and cleavage are lower. However, it tolerates mismatches in regions distant from the PAM sequence (TTTN or TTN) in the protospacer, and off-target cleavage issues may become more problematic when Cpf1 activity is improved for therapeutic purposes. In our study, we investigated off-target cleavage by Cpf1 and modified the Cpf1 (cr)RNA to address the off-target cleavage issue. We developed a CRISPR-Cpf1 that can induce mutations in target DNA sequences in a highly specific and effective manner by partially substituting the (cr)RNA with DNA to change the energy potential of base pairing to the target DNA. A model to explain how chimeric (cr)RNA guided CRISPR-Cpf1 and SpCas9 nickase effectively work in the intracellular genome is suggested. In our results, CRISPR-Cpf1 induces less off-target mutations at the cell level, when chimeric DNA-RNA guide was used for genome editing. This study has a potential for therapeutic applications in incurable diseases caused by genetic mutation.

## Background

The CRISPR-Cas system is a bacterial immune system and it is now widely applied to various organisms for target-specific genome editing[1]. One of the CRISPR system, CRISPR-Cas9 RNA-guided endonuclease is widely used to specifically correct or control genes of interest based on its ability to cut double-stranded DNA [1–5]. The other component, CRISPR-Cas12a (Cpf1) is lately reported and it is a class II, type V effector nuclease that has a bi-lobed structure composed of nuclease and recognition domains, similar to Cas9 [6, 7]. Cpf1 binds the target DNA helix via a single-stranded (cr)RNA, forming a DNA-RNA hybrid duplex [6, 8]. Cpf1 has attracted attention as an excellent genome-engineering tool because it overcomes certain limitations of Cas9 [9–12]. In contrast to Cas9, which recognizes guanine (G)-rich sequences, Cpf1 recognizes specific thymine (T)-rich protospacer adjacent motif (PAM) sequences (TTTN or TTN) and can specifically induce double-strand DNA cleavage [7, 13]. Nowadays, CRISPR-Cpf1 is broadly applicable, from microorganisms to humans, and thus, many researchers have been tried to engineer the CRISPR-Cpf1 protein or (cr)RNA for specific purpose [14–17]. In particular, for safety issues, (cr)RNAs, which are more accessible than proteins, can be engineered to improve the genome-editing specificity. Cpf1 recognizes a protospacer sequence of 24 bases, including a 5-10-base seed sequence known as the PAM sequence [18]. CRISPR-Cpf1 is more sensitive to mismatches between the target DNA and the gRNA than CRISPR-Cas9 is; when a mismatch is introduced into the seed sequence in the protospacer, its cleavage activity is significantly inhibited [19, 20]. However, there is still a possibility of mismatch cleavage in regions other than the seed region, which can currently not be detected and thus, the off-target cleavage issue is not entirely resolved. Substitution of the (cr)RNA with DNA possibly increase the target specificity by changing the binding energy between the guide and target [21]. In addition, DNA is more stable than RNA in aqueous solution and thus is more easily applicable in genome editing and more convenient for product commercialization. Therefore, in the current study, we partially replaced the gRNA for CRISPR-Cpf1 with DNA in an attempt to improve the accuracy of the system and to diversify the conventional CRISPR-Cpf1 genome editing tool. Chimeric DNA-RNA guides with high target specificity were screened by measuring the cleavage efficiency of each chimeric guide-Cpf1 complexes based on on- and off-target DNA sequences. We thus identified a chimeric guide with high accuracy, without on-target cleavage compensation. This novel system is advantageous in terms of safety and has application potential for various purposes *in vivo* [22, 23], and will eventually be useful for gene therapy for diseases caused by genetic defects.

## Results

### Effects of DNA substitution in the 5’, 3’-end of the (cr)RNA on Cpf1 activity

The (cr)RNA and target DNA recognition site of Cpf1 have been well characterized [6, 8]. Cpf1 proteins, including *Acidaminococcus* sp. Cpf1 (AsCpf1), generally bind with the target DNA to form a target DNA-gRNA duplex in a non-target-sequence-dependent fashion via positively charged amino acids in the REC1, REC2, and RuvC domains. In particular, the seed proximal region (5-10 bp from the PAM) is recognized by the amino acid residues of WED-REC1-RuvC and is essential for target DNA recognition, and it constitutes a protospacer with relatively less sensitive PAM distal regions [6]. Cpf1 amino acid residues that recognize the 2’-OH group in the (cr)RNA are known (His872, Glu786, Asn175, Arg176, Arg192, Arg518, Asn515, Gly270, Lys273, Gln286), and the 3’-end of the gRNA is denatured by tryptophan, which is structurally universal, endowing important properties for CRISPR-Cas12a activity. To change the interaction between the (cr)RNA and Cpf1 protein, we replaced part of the (cr)RNA sequence in the 5’-side hairpin region, a single-strand region separated by a tryptophan residue (Y382) at the 3’-side and the protospacer DNA-RNA hybrid region, with DNA **(Figure 1A)**. First, we sequentially replaced 4-nt regions of the RNA guide with DNA starting from the 3’ end, the cleavage efficiency for each target (*DNMT1, CCR5*) was compared **(Figure 1B, Supplementary Figures S1, S2)**. PCR amplicon cleavage of the two target genes revealed that up to 8 nt of DNA substitution starting from the 3’-end did not significantly affect the target cleavage efficiency. However, when more than 9 consecutive bases in the guide were substituted, this resulted in a significant decrease in on-target sequence cleavage efficiency, and no cleavage was observed when more than 12 nt were substituted in Cpf1 from various sources **(Supplementary Figure S3)**. This suggests that 2’-OH recognition of the (cr)RNA is largely conserved among Cpf1 proteins and that 2’-OH recognition by CRISPR-Cas12a increases from the PAM (TTTN or TTN) distal to the PAM proximal region. This is in line with previous findings in experiments in which mismatches were sequentially introduced into the gRNA and target DNA heteroduplex regions [18]. When the RNA nucleotides from the 5’-end of the hairpin region other than the protospacer were replaced with DNA, the chimeric DNA-RNA-guided Cpf1 showed a significant decrease in cleavage efficiency when compared to Cpf1 with the wild-type guide **(Figure 1C)**. Similarly, Cpf1 activity decreased when the conserved region of the 5’-hairpin structure was replaced or removed [7]. These findings indicate that the 2’-OH in the 5’-hairpin region, which forms a pseudoknot, is critically required for Cpf1 protein recognition of the (cr)RNA.

**Figure 1.**
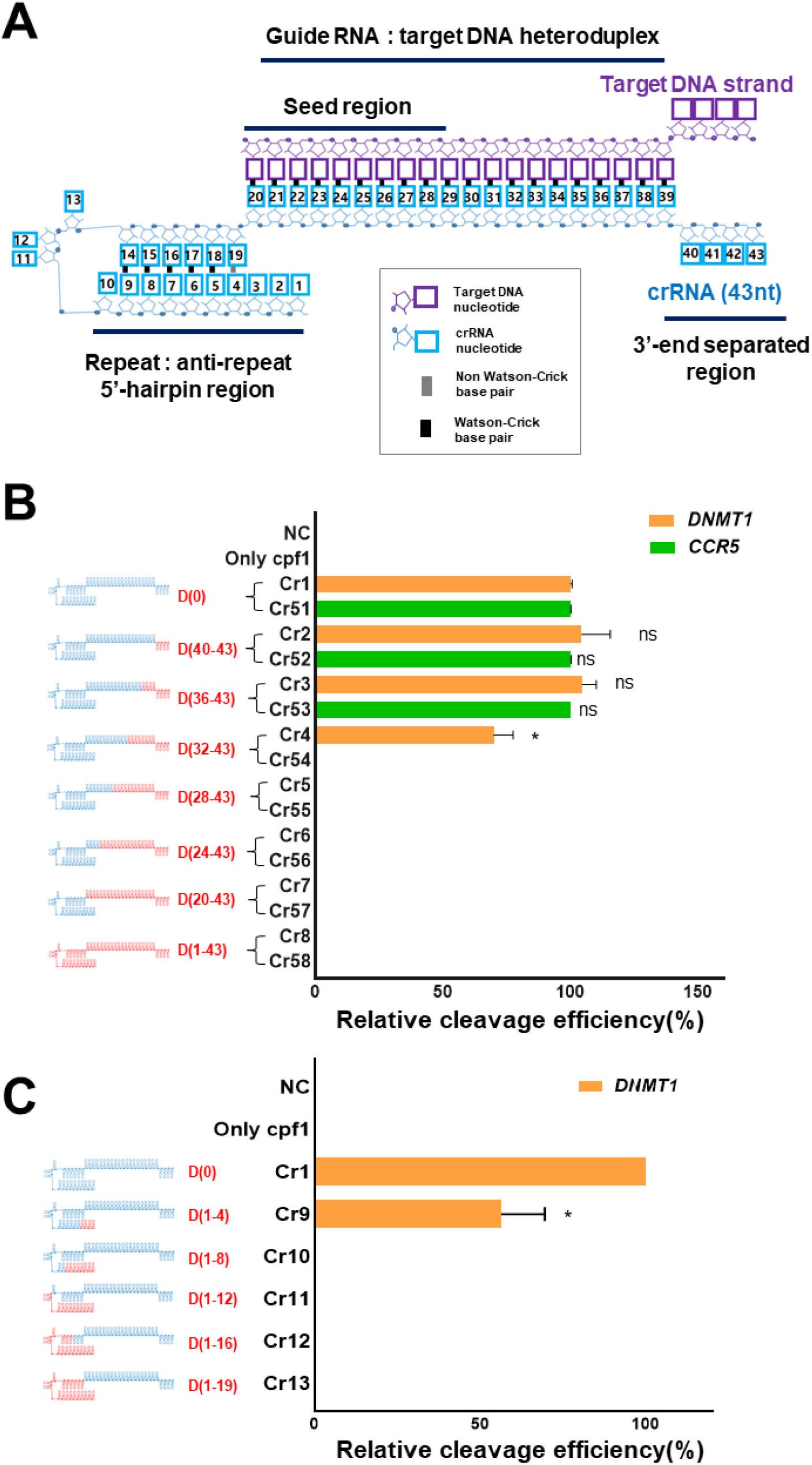
Target DNA cleavage by CRISPR-Cas12a (Cpf1) using chimeric DNA-RNA guides. **(A)** Schematic representation of interactions of AsCpf1 (cr)RNA (colored in cyon) with target DNA (colored in purple) [6]. The AsCpf1 (cr)RNA was numbered from 5’-end to 3’-end. **(B)** The target DNA amplicon cleavage efficiency of AsCpf1 with partial DNA substitution of (cr)RNA was determined. The (cr)RNA was replaced with DNA from 3’-end with 4nt interval. The RNA portion of the (cr)RNA is shown in blue, and the DNA portion is shown in red, respectively (‘D’ indicates a DNA and the number of substituted DNA nucleotides is indicated). X axis indicates for efficiency of the target gene (*DNMT1* (orange), *CCR5* (green)) cleavage by AsCpf1 using various chimeric DNA-RNA guides (DNA substitution of 4-nt from the 3′-end of the (cr)RNA). Y axis indicates for used chimeric (cr)RNAs in cleavage experiment. **(C)** Target gene (*DNMT1* (orange)) cleavage by AsCpf1 using chimeric DNA-RNA guides (Serial 4-nt DNA substitution from the 5′-end of the (cr)RNA). All the cleavage efficiency was calculated from agarose gel separated band intensity (cleaved fragment intensity (%) / total fragment intensity (%)) and normalized to wild-type (cr)RNA (crRNA 1 or crRNA 51). Data are shown as means ± s.e.m. from three independent experiments. *P*-values are calculated using a two-tailed Student’s t-test (ns: not significant, P*:<0.05, P**:<0.01, P***:<0.001, P****:<0.0001).

### Effects of DNA substitution in the seed region of the (cr)RNA on Cpf1 activity

Next, we replaced single nucleotides in the seed region of the guide, which is located close to the PAM sequence and required for target DNA-gRNA heteroduplex formation, with DNA bases and assessed the effects on Cpf1 protein activity. Target DNA cleavage experiments were performed for two genes (*DNMT1, CCR5*) **(Figure 2)**. Target gene cleavage efficiency was reduced after DNA replacement in the seed region of (cr)RNA up to 7-nt (crRNA17-23) from the PAM for target sequence of *DNMT1* **(Figure 2A)** and 5-nt distance region from the PAM (crRNA65) for *CCR5* **(Figure 2B)** gene respectively. In addition, according to the target sequences in *DNMT1* and *CCR5*, different pattern of the cleavage efficiencies were obtained by replacing the seed-region RNA nucleotides with DNA nucleotides. Thus, the recognition of 2’-OH in the (cr)RNA depends on the nucleotide sequence of the target gene. Next we evaluated the effects of gradual DNA substitutions in groups of four nucleotides in the seed region (crRNA14-16). As expected, this led to a greater reduction in target cleavage efficiency than single base substitution **(Figure 2A, top)**. Taken together, these findings suggested that DNA substitution in the 3’ region (up to 8 nt) better maintained the cleavage activity than DNA substitution in the seed or 5’-hairpin region of the guide. All the chimeric (cr)RNA forms evaluated were screened for Cpf1 activity in an additional cleavage efficiency test.

**Figure 2.**
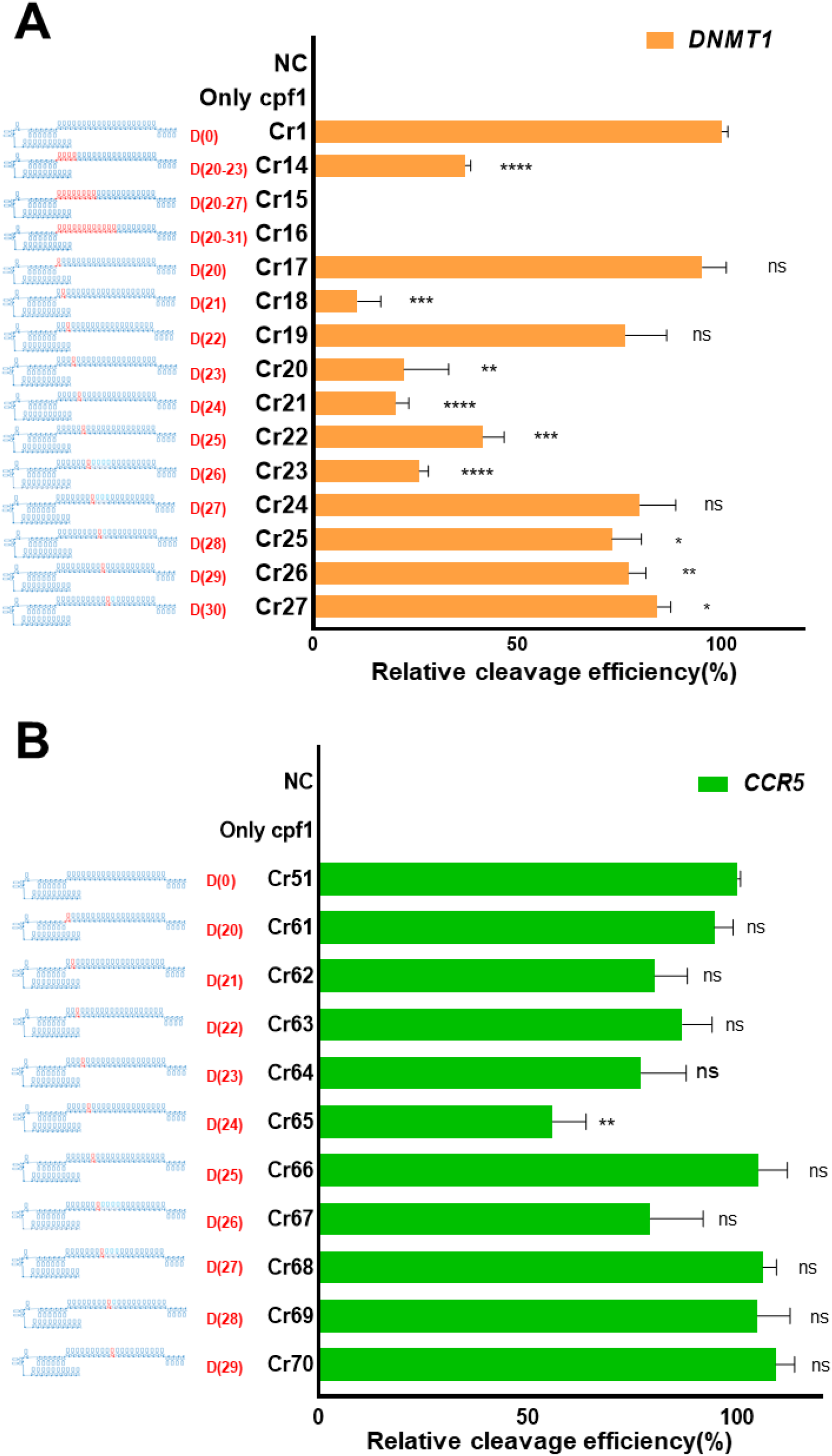
Target DNA cleavage by Cpf1 using a chimeric DNA-RNA guide in which the seed region of the (cr)RNA is replaced with DNA. **(A)** Comparative analysis of the target (*DNMT1*: orange) DNA cleavage efficiency using various (cr)RNAs harboring serial multiple or single DNA substitutions in the seed region close to the PAM (TTTN). **(B)** Comparative analysis of the target (*CCR5*: green) DNA cleavage efficiency. AsCpf1 (cr)RNA was serially replaced with single DNA nucleotides from the PAM. The RNA portion of the (cr)RNA is shown in blue, and the DNA portion is shown in red (‘D’ indicates a DNA and the number of substituted DNA nucleotides is indicated). All the cleavage efficiency was calculated from agarose gel separated band intensity (cleaved fragment intensity (%) / total fragment intensity (%)) and normalized to wild-type (cr)RNA (crRNA 1 or crRNA 51). Data are shown as means ± s.e.m. from three independent experiments. *P*-values are calculated using a two-tailed Student’s t-test (ns: not significant, P*:<0.05, P**:<0.01, P***:<0.001, P****:<0.0001).

### Target DNA cleavage specificity of chimeric DNA-RNA-guided Cpf1

To confirm the target DNA specificity after DNA substitution of the AsCpf1 guide, the on- and off-target cleavage efficiencies for various genes (*DNMT1, FANCF, GRIN2B, EMX1*) were measured **(Figure 3)**. Off-target sequences for each target sequence with up to 7 mismatches were predicted *in silico* using Cas-OFFinder [24] **(Supplementary Table S1)**, so that highly probable off-target cleavage was interrogated **(Figure 3, inset table)**, and the on/off cleavage ratio was calculated to compare the target specificity **(Figure 4)**. First, we measured on- and off-target cleavage efficiencies of the chimeric (cr)RNAs presented in **(Figures 1, 2)** for *DNMT1*, which revealed that three off-target sites showed different cleavage efficiency relative to the on-target sequence **(Figure 3A, B)**. In this assay, we used unsaturated conditions for on-target cleavage and saturated conditions for off-target cleavage not to misinterpret the cleavage efficiency. 3’-end substitution of nucleotides +8 and +12 in the (cr)RNA resulted in a significant decrease in cleavage at off-target sites 1 and 2, and no cleavage was observed for off-target site 3 **(Figure 3A)**. In the case of 3’-end 8-nt continuous DNA substitution of the (cr)RNA, the specificity increased by 10.22-fold, without compensation of on-target cleavage efficiency **(Figure 4A)**. Increased target specificity was also observed for the FnCpf1 effector **(Supplementary Figure S4)**. For *DNMT1* target and off-target amplicons, FnCpf1 showed robust on-target cleavage, but almost no off-target cleavage when a 3’-end 8-nt DNA-substituted (cr)RNA was used. In the case of single-nucleotide DNA substitution in the (cr)RNA seed region, up to 11 gradual single substitutions from the PAM sequence showed high correlation between off-target and on-target cleavage sites **(Figure 3B)**. In particular, one-by-one single-nucleotide DNA substitution in the seed region tended to significantly increase on-target cleavage compensation, despite the increase in target specificity due to the absence of off-target cleavage. As a result, there was a significant increase in cleavage specificity (0.11 ~4.15 fold) for chimeric guides ((cr)RNA No. 14, 20, 21, 22, 23) that did not induce DNA off-target cleavage **(Figure 4B)**. For the *DNMT1* target gene, consecutive DNA substitutions from the 3’-end of the guide to the eighth nucleotide resulted in less on-target compensation and higher target cleavage specificity than DNA substitutions in the seed region close to the PAM sequence **(Figures 4A, 4B)**. In further experiments, we used RNA-DNA chimeric guides with 3’-end substituted sequences to determine the target specificity for more gene sequences **(Figures 3C-E, 4C-E)**. When 8 nucleotides were substituted at the 3’-end of the guide, on-target cleavage efficiency was maintained or slightly increased for three genes (*FANCF, GRIN2B*, and *EMX1*), and off-target cleavage was decreased relative to that of the wild-type (cr)RNA at off-target site 1 in *FANCF* and off-target site 2 in *GRIN2B* **(Figure 3 C-E)**. At other off-target sites for three target genes, mismatches in the seed region inhibited cleavage, despite the use of a high concentration of Cpf1-guide complex, which induced perfect on-target cleavage **(Supplementary Figure S1)**. *In-vitro* cleavage experiments showed similar results in that the specificity (on-/off-target cleavage ratio) was substantially increased (0.88~2.07 fold) after 3’-end (up to 8-nt) DNA substitution of the guides for the three additional genes **(Figure 4C-E)**. These results showed that 3’-end DNA substitution of the RNA guide increases target DNA cleavage specificity without significant effect of the nucleotide sequence itself.

**Figure 3.**
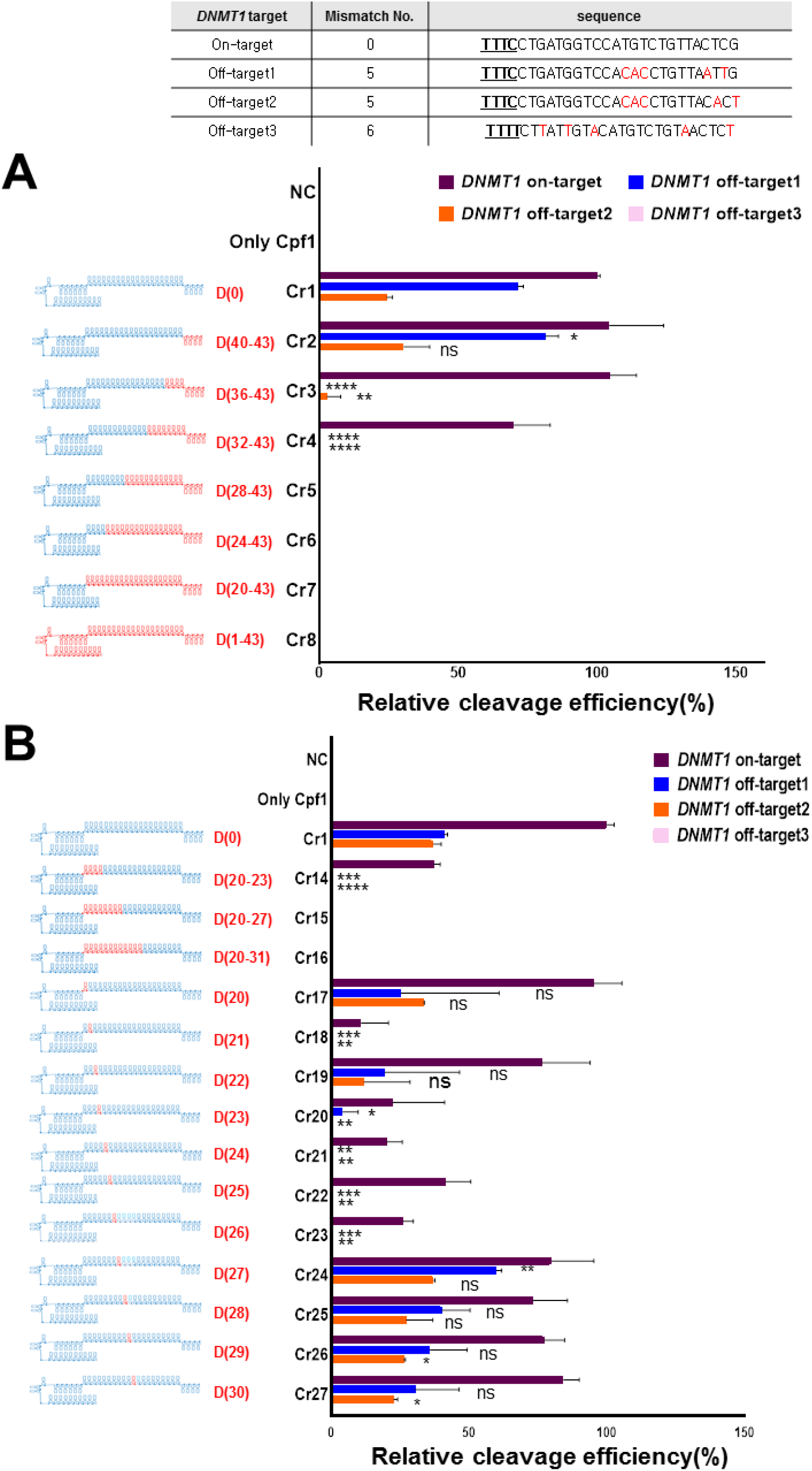
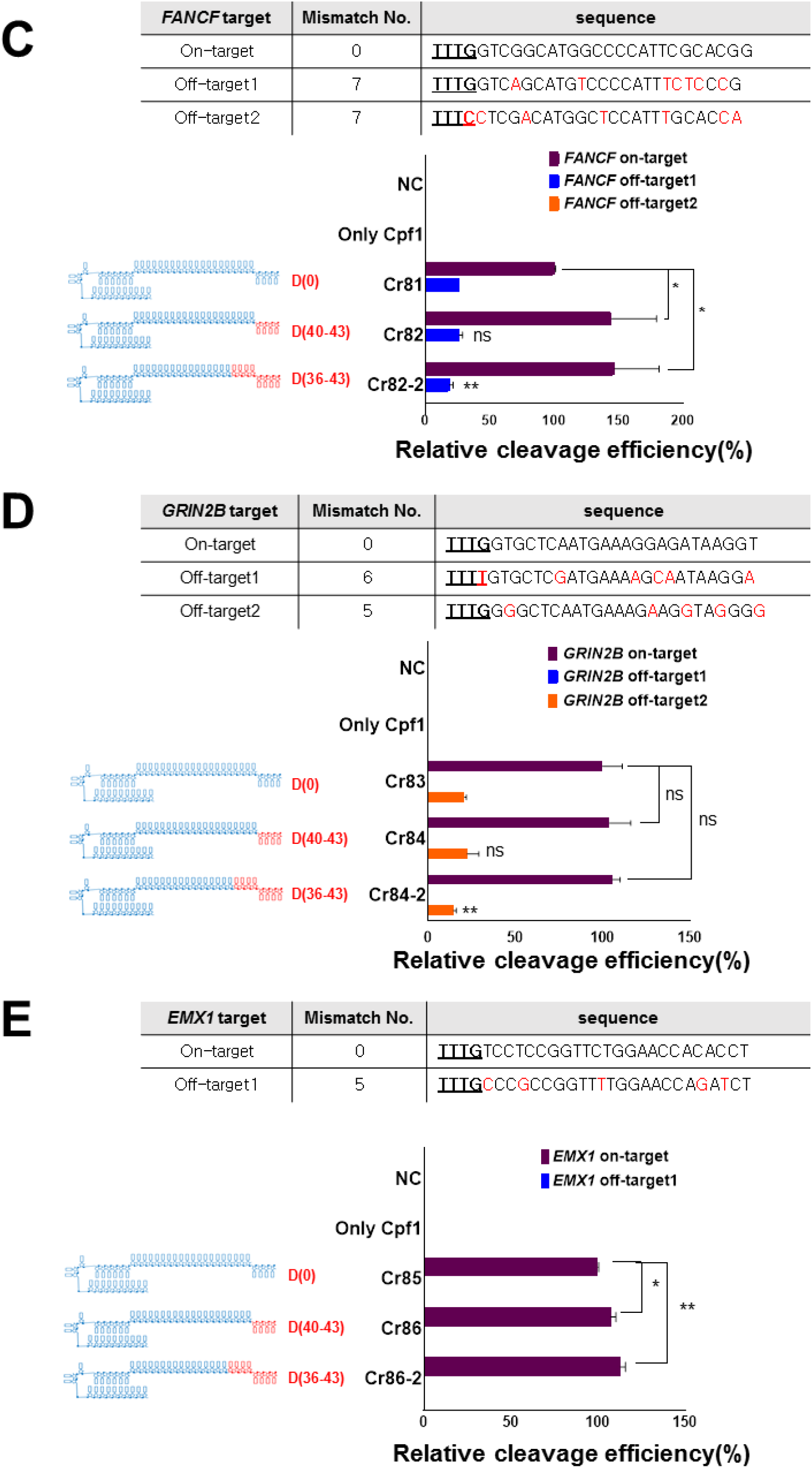
Off-target cleavage assay of Cpf1 using chimeric DNA-RNA guides. **(A)** Target and off-target *DNMT1* DNA cleavage experiments to confirm the target specificity of AsCpf1 using a chimeric DNA-RNA guide (serial 4-nt DNA substitutions from the 3’-end; on-target (dark brown), off-target 1 (blue), off-target 2 (orange), off-target 3 (pink)). The RNA portion of the Cpf1 (cr)RNA is shown in blue, and the DNA portion is shown in red (‘D’ indicates a DNA and the number of substituted DNA nucleotides is indicated). **(B)** Target and off-target *DNMT1* DNA cleavage experiments to confirm the target specificity of AsCpf1 using a chimeric DNA-RNA guide (DNA substitutions in the seed region of (cr)RNA; on-target (dark brown), off-target 1 (blue), off-target 2 (orange), off-target 3 (pink)). **(C)** Relative cleavage efficiency of target (dark brown) and off-target (off1: blue, off2: orange) of *FANCF* DNA using chimeric DNA-RNA guided (3’-end 4-bp, 8-bp DNA substitutions) AsCpf1. **(D)** Relative cleavage efficiency of target (dark brown) and off-target (off1: blue, off2: orange) of *GRIN2B* DNA using chimeric DNA-RNA guided (3’-end 4-bp, 8-bp DNA substitutions) AsCpf1. **(E)** Relative cleavage efficiency of target (dark brown) and off-target (off1: blue) of *EMX1* DNA using chimeric DNA-RNA guided (3’-end 4-bp, 8-bp DNA substitutions) AsCpf1. All the cleavage efficiency was calculated from agarose gel separated band intensity (cleaved fragment intensity (%) / total fragment intensity (%)) and normalized to wild-type (cr)RNA (crRNA 1 for *DNMT1*, crRNA 81 for *FANCF*, crRNA 83 for *GRIN2B*, crRNA 85 for *EMX1*). Data are shown as means ± s.e.m. from three independent experiments. *P*-values are calculated using a two-tailed Student’s t-test (ns: not significant, P*:<0.05, P**:<0.01, P***:<0.001, P****:<0.0001). On- and off-target sequence information for each gene is shown at the top. The PAM sequence is underlined and shown in bold. Mismatch sequences to wild-type reference are shown in red.

**Figure 4.**
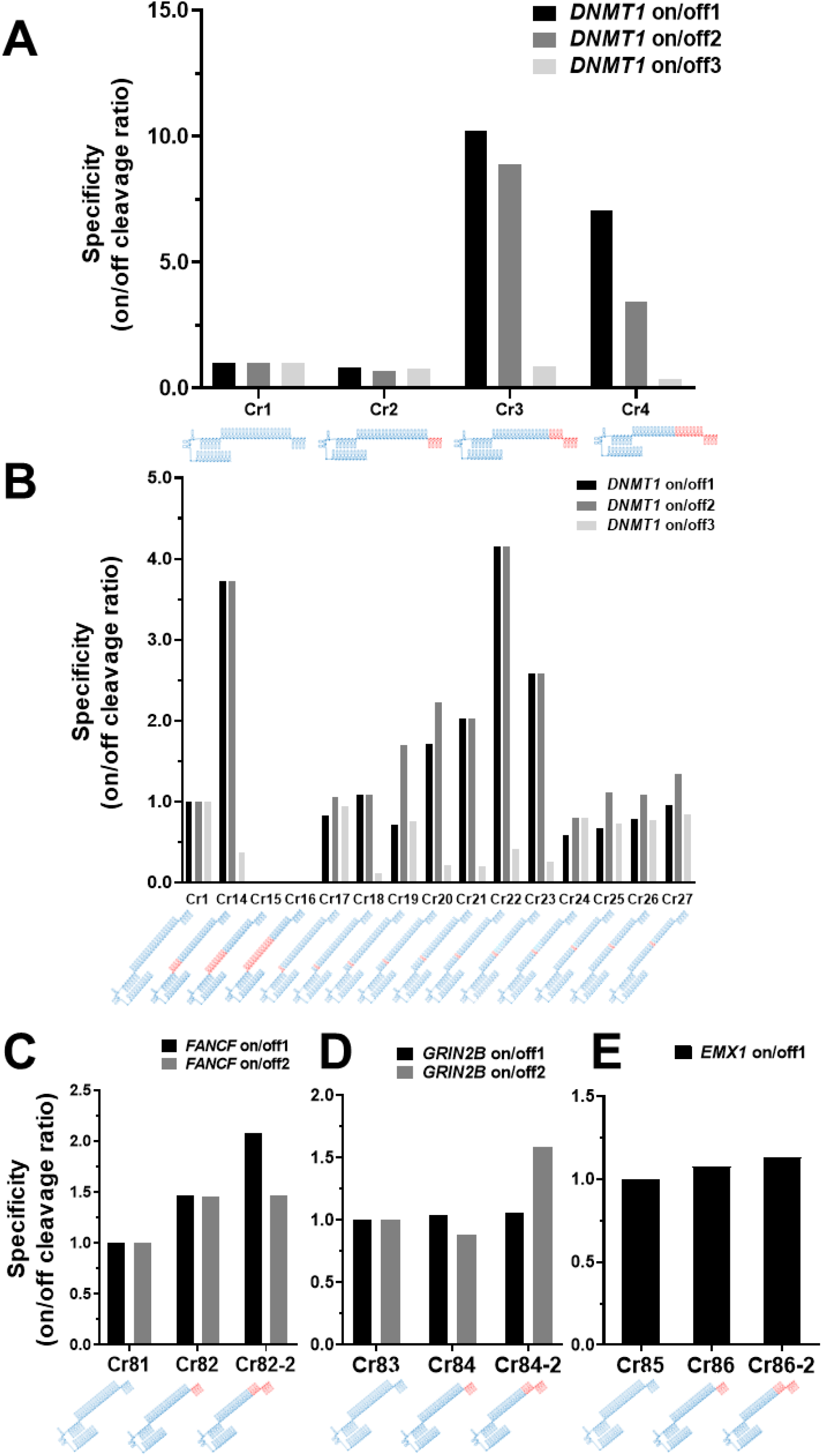
Comparison of target specificity of chimeric DNA-RNA guides. **(A)** Comparison of the *DNMT1* target specificity (on/off cleavage ratio (%)) of AsCpf1 using chimeric DNA-RNA guides with serial 4-nt DNA substitutions from the 3’-end of the (cr)RNA (Substituted DNA nucleotides in (cr)RNA is shown in red color and RNA is shown in blue color, respectively). Target specificity was calculated from the results in **(Figure 3)** by dividing on-target cleavage efficiency by off-target cleavage efficiency. Target specificity is shown in black (on/off 1), dark gray (on/off 2), and light gray (on/off 3). **(B)** Comparison of the *DNMT1* target specificity of AsCpf1 using chimeric DNA-RNA guides with DNA substitutions in the seed region of the (cr)RNA. Target specificity is shown in black (on/off 1), dark gray (on/off 2) and light gray (on/off 3). **(C-E)** Comparison of *FANCF, GRIN2B*, and *EMX1* target specificities of AsCpf1 using chimeric DNA-RNA guides with serial 4-nt DNA substitutions from the 3’-end of the (cr)RNA). Target specificity is shown in black (on/off 1) and gray (on/off2). The number of substituted DNA nucleotides in AsCpf1 (cr)RNA used for each gene targeting is indicated by red color.

### 3’-End DNA-substituted chimeric DNA-RNA-guided Cpf1 shows versatile activity in human cells

For the *in-vivo* application of Cpf1 using various chimeric RNA-DNA guides with high target cleavage specificity, we confirmed whether target-specific genome editing can be induced at the cellular level **(Figure 5)**. Indels in *DNMT1* were induced by delivering various chimeric DNA-RNA guides and purified Cpf1 proteins into HEK293FT cells. We observed that all chimeric guides induced mutations in the target gene **(Figure 5A-C)**, mostly deletions **(Figure 5D, Supplementary Figure S5)**. In this cell system, 3’-end DNA substitution of the (cr)RNA led to a sharp decrease in indel formation efficiency at the target sequence **(Figure 5A)**, unlike the findings in the *in-vitro* PCR amplicon cleavage experiment **(Figure 1B)**. In contrast, when a seed-region DNA-substituted guide was used, indels were induced at an efficiency similar to that of *in-vitro* amplicon cleavage **(Figure 5B)**. In particular, the CRISPR-Cpf1 activity in cells was decreased when the DNA substitution was located in the seed region rather than in the PAM distal region **(Figure 5A, B)**. In case of DNA substitution at the 5’-end of the guide, the indel formation rate was reduced to a level similar to that the *in-vitro* cleavage experiment **(Figure 1C)**. This result indicated that Cpf1 is highly deactivated upon alteration in the conserved 5’-end structure of (cr)RNA **(Figure 5C)**. Together, these results suggested that the difference in cleavage activity between PCR amplicons and endogenous intracellular loci might be related to various effects of the intracellular condition, such as degradation issues by DNA exonuclease [25], structural hindrances by binding of histones or other proteins [26], or the supercoiled condition of intracellular chromosomes [27], which might affect the binding of CRISPR effector proteins *in vivo* [28].

**Figure 5.**
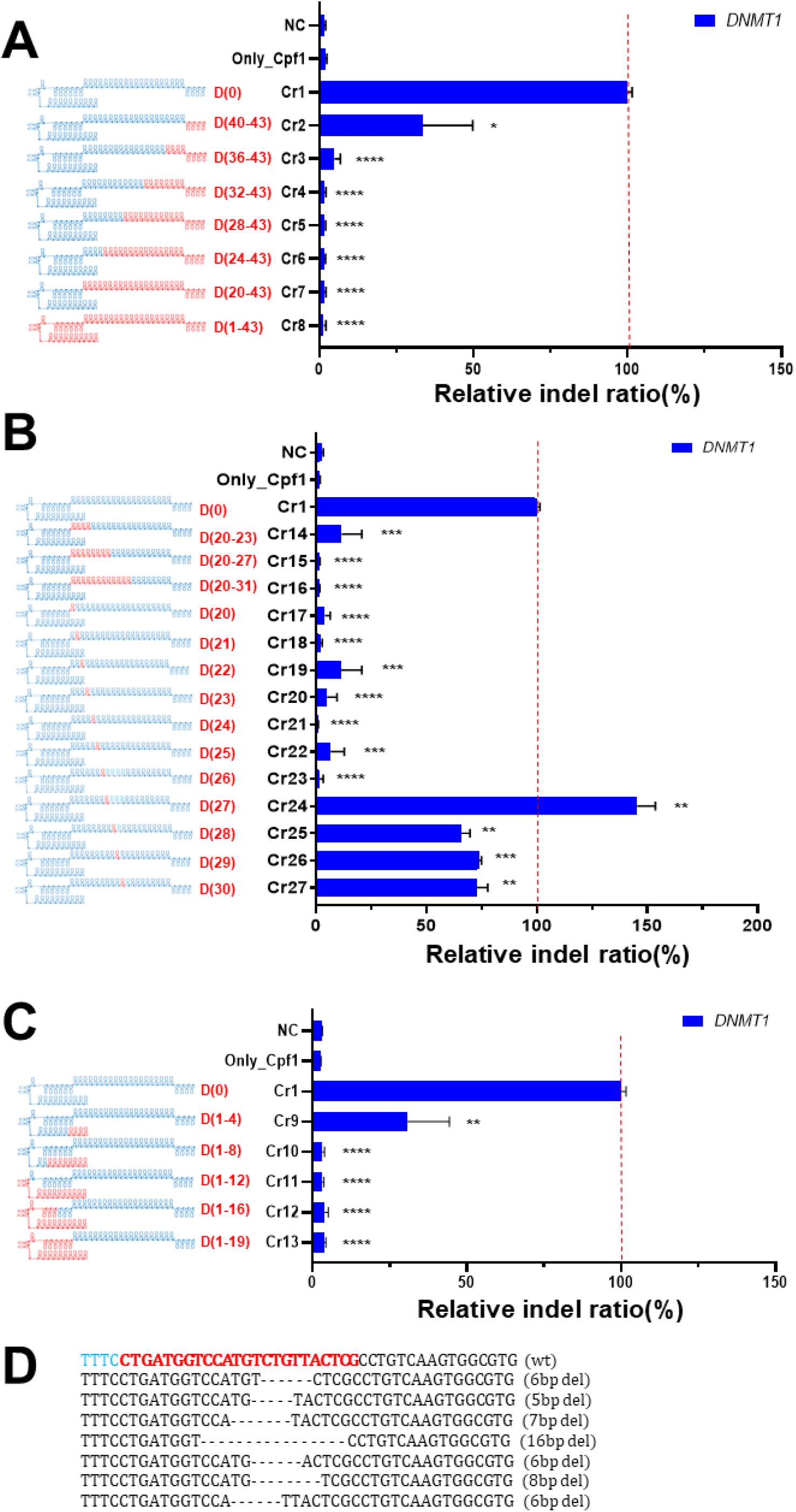
Intracellular genome editing using Cpf1 and chimeric DNA-RNA guides. **(A)** Endogenous locus (*DNMT1*) editing in HEK293FT cells using chimeric DNA-RNA guides with serial 4-nt DNA substitutions from the 3′-end of the (cr)RNA. The RNA portion of the Cpf1 (cr)RNA is shown in blue, and the DNA portion is shown in red (‘D’ indicates a DNA and the number of substituted DNA nucleotides is indicated). **(B)** Endogenous locus (*DNMT1*) editing in HEK293FT cells using chimeric DNA-RNA guides with DNA substitutions in the seed region of the (cr)RNA. **(C)** Endogenous locus (*DNMT1*) editing in HEK293FT cells using chimeric DNA-RNA guides with serial 4-nt DNA substitutions from the 5′-end of the (cr)RNA. **(D)** Representative genome editing pattern of AsCpf1 using chimeric DNA-RNA guides. Sequencing data is obtained from *DNMT1* targeted amplicon sequencing. Deleted base is shown by dashed line. PAM sequence (TTTN) and target sequence for AsCpf1 is shown in cyon and red color, respectively. All the relative indel ratio was calculated from targeted amplicon sequencing (indel frequency (%) = mutant DNA read number / total DNA read number) and normalized to wild-type (cr)RNA (crRNA 1 for *DNMT1*). Data are shown as means ± s.e.m. from three independent experiments. *P*-values are calculated using a two-tailed Student’s t-test (ns: not significant, P*:<0.05, P**:<0.01, P***:<0.001, P****:<0.0001).

### 3’-End modification of the chimeric DNA-RNA recovers Cpf1 activity in human cells

To determine and prevent the degradation of the chimeric guide by 3′-end DNA exonuclease [29], we performed target-specific genome editing using a chimeric guide of which the 3′ end (with 4-nt or 8-nt DNA substitution) moiety was chemically modified with phosphorothioate (PS) **(Figure 6A, Supplementary Table S1)**. To test AsCpf1 activity when using 3′-end-modified chimeric guides, the *DNMT1* target cleavage was assessed *in vitro* **(Figure 6A)**. The target DNA cleavage efficiency of the 3’-end-modified guide was similar to that of the wild-type gRNA, without on-target cleavage compensation. On the other hand, upon 3′-end 8-nt DNA substitution of the guide, cleavage at off-target sites 1 and 2 was reduced or not detected, respectively **(Figure 6A)**. Thus, the target specificity was increased (1.03~10.35fold) as compared to that of wild-type or 3′-end +4 DNA-substituted guides, even with 3′-end modification **(Figure 6B)**. To evaluate Cpf1 activity with chemically modified chimeric guides in mammalian cells, 3’-end modified chimeric guides and purified Cpf1 proteins were delivered together into HEK293FT cells to induce indels in the target gene **(Figure 6C, Supplementary Figure S6)**. The decrease in on-target cleavage **(Figure 5A)** was restored by 2.3-fold when compared to that of non-chemically modified guide with 4-nt DNA substitution **(Figure 6C)**. However, in the case of 8-nt DNA substitution, the editing efficiency was not enhanced by 3′-end PS modification. These results suggested that Cpf1 with 3′-end 4-nt DNA chimeric guides, for which part of the (cr)RNA is separated from the target DNA strand, can be effectively applied in cells through extension of the lifetime of the (cr)RNA. For 3′-end 8-nt DNA chimeric (cr)RNA, which has both base-paired and separated regions to target DNA strand, further improvement other than 3′-end chemical modification is needed to activate the Cpf1 on endogenous loci inside the cell.

**Figure 6.**
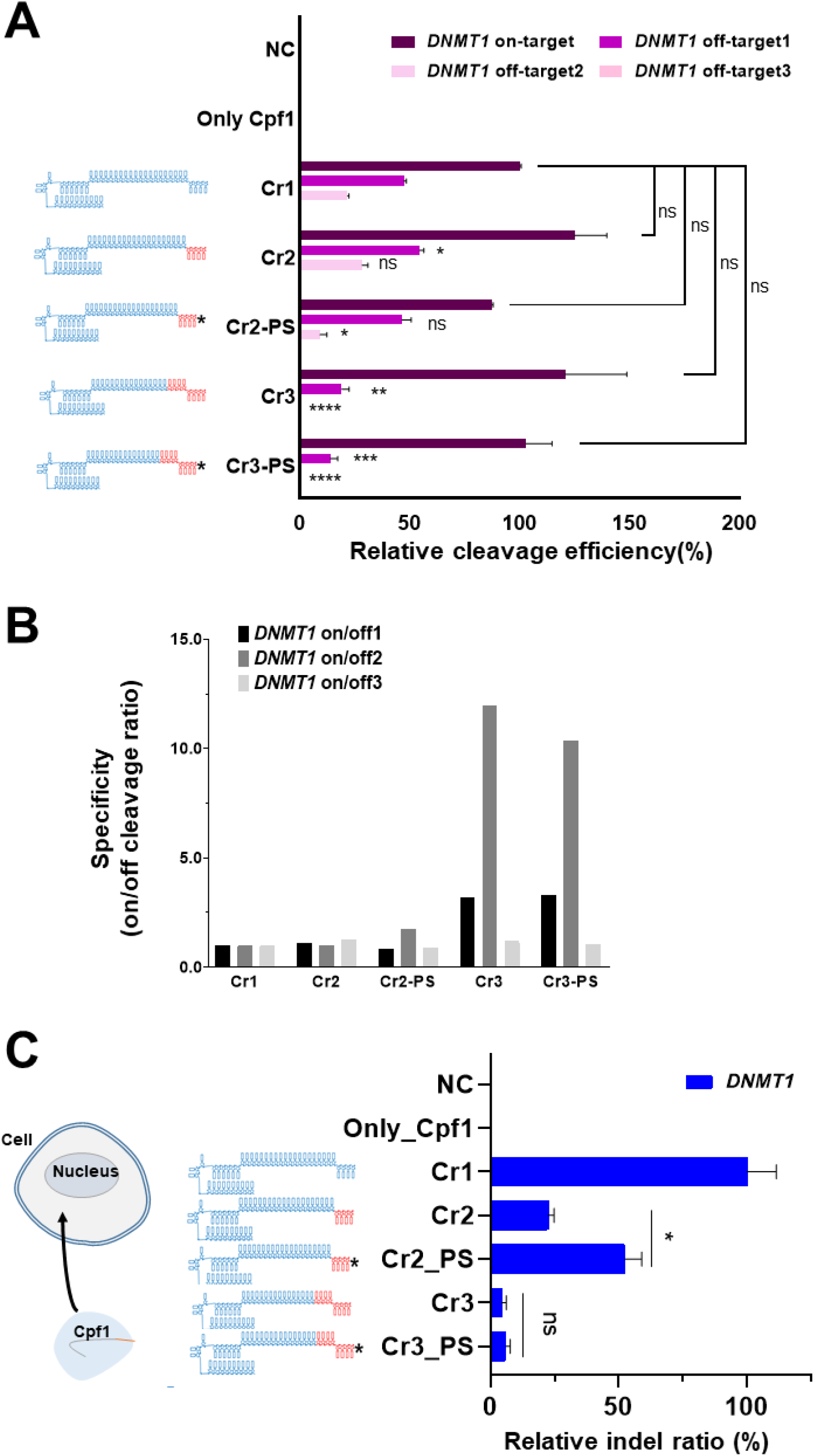
Intracellular genome editing of Cpf1 using 3′-end chemically modified chimeric DNA-RNA guides. **(A)** *DNMT1* on/off-target amplicon cleavage by AsCpf1 using chimeric DNA-RNA guides with 3’-end 4-nt or 8-nt DNA substitution and 3’-end PS modification. The relative cleavage efficiency (%) of on-target (dark brown), off-target 1 (dark pink), off-target 2 (light purple) and off-target 3 (light pink)) is indicated by different colors. The RNA portion of the Cpf1 (cr)RNA is shown in blue, and the substituted DNA portion is shown in red. Asterisk indicates the 3’-end PS modification of (cr)RNA. All the cleavage efficiency was calculated from agarose gel separated band intensity (cleavage efficiency (%) = cleaved fragment intensity / total fragment intensity) and normalized to wild-type (cr)RNA (crRNA 1 for *DNMT1*). Data are shown as means ± s.e.m. from three independent experiments. *P*-values are calculated using a two-tailed Student’s t-test (ns: not significant, P*:<0.05, P**:<0.01, P***:<0.001, P****:<0.0001). **(B)** Determination of target specificity (on/off cleavage ratio (%)) calculated from (A) of the chimeric DNA-RNA guide with 3′-end PS modification. Target specificity is shown in black (On/Off 1), dark gray (On/Off 2), and light gray (On/Off 3), respectively. **(C)** Intracellular *DNMT1* editing using 3′-end PS-modified chimeric DNA-RNA guides. Relative indel ratio (%) is calculated by targeted amplicon sequencing from *DNMT1* site in HEK293FT cells (indel frequency (%) = mutant DNA read number / total DNA read number) and normalized to wild-type (cr)RNA (crRNA 1 for *DNMT1*). Data are shown as means ± s.e.m. from three independent experiments. *P*-values are calculated using a two-tailed Student’s t-test (ns: not significant, P*:<0.05, P**:<0.01, P***:<0.001, P****:<0.0001).

### SpCas9 nickase in combination with chimeric DNA-RNA-guided Cpf1 ensures highly specific genome editing in intracellular conditions

To improve the genome editing efficiency of (cr)RNA with 3′-end 8-nt DNA substitution, we tried to address the supercoiling issue in intracellular conditions, which leads to highly compact genomic DNA. Based on a previous study, we attempted to reduce or eliminate supercoiling at target sequences using a dead or nickase-type CRISPR system [28]. To this end, we confirmed whether the genomeediting efficiency of Cpf1 using a 3′-end 4- or 8-nt DNA-substituted (cr)RNA could be improved by using it in combination with dead SpCas9 or SpCas9 nickase **(Figure 7)**. A single-guide RNA was designed **(Supplementary Figure S7A, Table S3)**for the binding of dead (D10A, H840A) or nickase (D10A) SpCas9 16-bp away from the *DNMT1* sequence, and purified SpCas9-sgRNA complex was transfected into cells together with Cpf1 protein. Interestingly, compared to non-SpCas9-treated cells, dead-type SpCas9-treated cells showed no significant change in target genome indel frequency (%) upon 4- or 8-nt DNA substitution of the (cr)RNA, but treatment with nickase-type SpCas9 was found to dramatically increase the indel frequency (%) in the target sequence **(Figure 7A, Supplementary Figure S7B)**. In particular, we observed a greater increase in indel ratio (%) of fold increase (22.1fold) for the 8-nt DNA-substituted (cr)RNA, which contains a region that base-pairs with the target DNA strand, than for the 3’-end 4-nt DNA substituted (cr)RNA **(Figure 7B)**. When only dead or nickase-type SpCas9 complex was delivered, no indels were generated. To test the synergistic effect of the chemical modification of the chimeric (cr)RNA and nickase activity, we delivered all the materials into the cell at the same time. But, nickase SpCas9 (D10A) and 3’-end modification of the (cr)RNA showed no synergistic effect on genome editing efficiency **(Supplementary Figure S8)**. These results suggested that Cpf1 efficiently operates in a chimeric (cr)RNA-dependent manner only when a nick is formed near the target DNA sequence under intracellular conditions. To investigate the off-target issues generated by combination of nickase and Cpf1 effector, we performed Guide-seq and targeted amplicon sequencing to analyze the off-target cleavage in a genome-wide **(Supplementary Figure S9A)**and target-specific manner **(Supplementary Figure S9B)**. As a result, when analyzed by Guide-seq, it was confirmed that off-target mutation at site 1 and 2 **(Figure 7C, Table)**for *DNMT1* target was not detected in the sample treated with nickase and chimeric (cr)RNA guided Cpf1 **(Figure 7C)**. Targeted amplicon sequencing was also performed for predicted off-target sites for the *DNMT1* target sequence as same with in-vitro cleavage assay **(Figure 3A, 3B)**, and no significant off-target indels were observed **(Supplementary Figure S9)**. However, when the plasmid target (*DNMT1*) was edited intracellularly, it was confirmed that the indel ratio (%) was significantly reduced at off-target 1, 2 **(Figure 7C, Supplementary Figure S10)** as in the *DNMT1* off-target cleavage experiment **(Figure 3A)**. As a result, when chimeric (cr)RNA was used intracellularly, it was confirmed that the specificity was significantly increased (63.34~167.94fold) **(Figure 7D)**. In conclusion, our data suggest that Cpf1 works effectively with chimeric DNA-RNA guides only in the absence of supercoiling, and nickase-chimeric guide combination ensures highly-specific genome editing with Cpf1 effector. On the basis of this concept, we suggest a model for the highly efficient and specific target DNA cleavage mechanism of Cpf1 that operates based on a chimeric DNA-RNA guide as shown in **Figure 8**.

**Figure 7.**
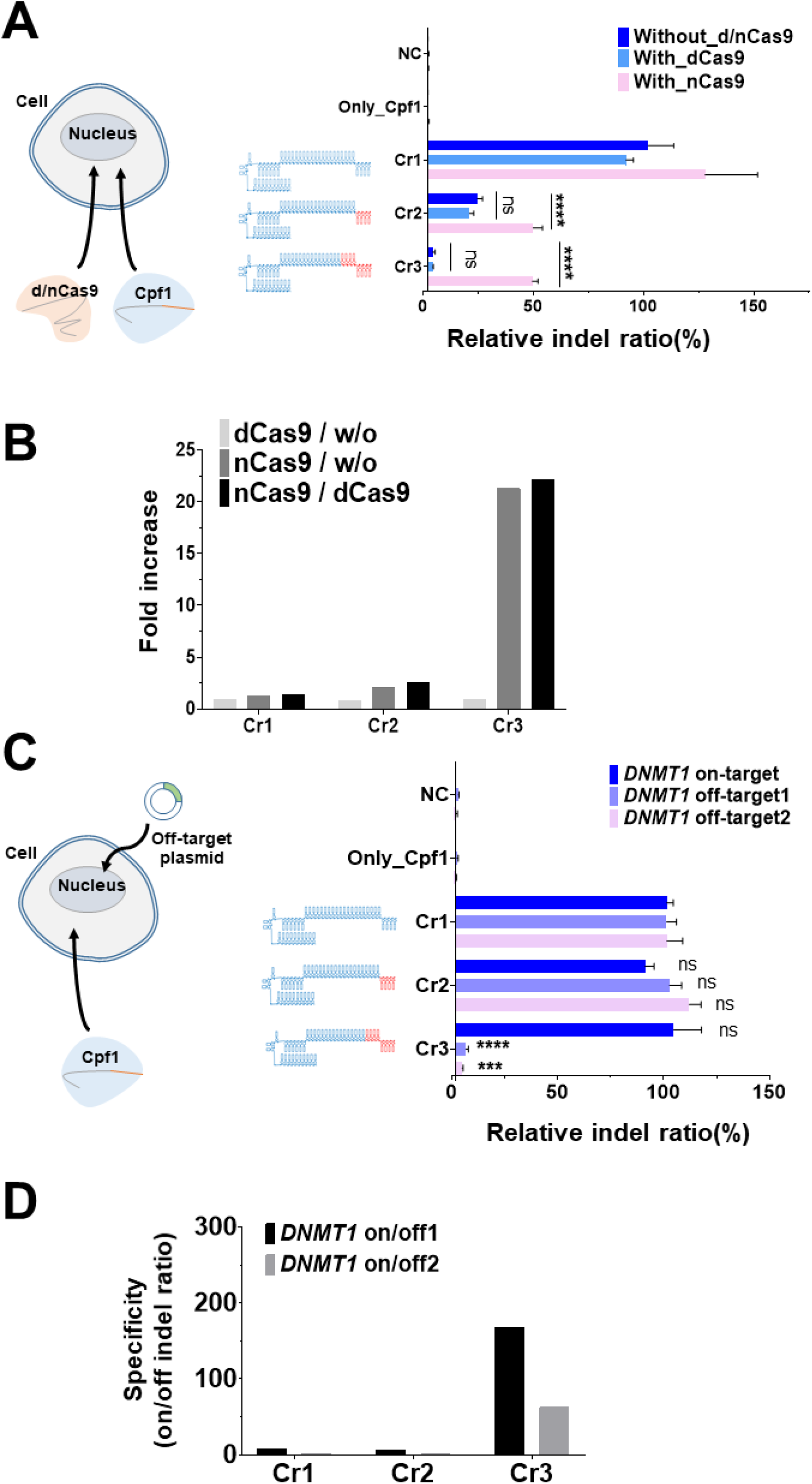
Cpf1 editing efficiency and specificity are enhanced by combination of a chimeric DNA-RNA guide and SpCas9 nickase. **(A)** Intracellular *DNMT1* gene editing in HEK293FT cell using 3′-end DNA-substituted chimeric DNA-RNA-guided Cpf1 and SpCas9 (D10A) nickase. The RNA portion of the Cpf1 (cr)RNA is shown in blue, and the substituted DNA portion is shown in red. X axis indicates the relative indel frequency (%). Only (cr)RNA - Cpf1 treated (dark blue) and combination with dCas9 (blue) or nCas9 (pink) treated samples were indicated, respectively. dCas9 and nCas9 indicates deactivated Cas9 (D10A, H840A) and nickase Cas9 (D10A), respectively. Indel ratio (%) is calculated by targeted amplicon sequencing from *DNMT1* site in HEK293FT cells (indel frequency (%) = mutant DNA read number / total DNA read number) and normalized to wild-type (cr)RNA (crRNA 1 for *DNMT1*). Data are shown as means ± s.e.m. from three independent experiments. *P*-values are calculated using a two-tailed Student’s t-test (ns: not significant, P*:<0.05, P**:<0.01, P***:<0.001, P****:<0.0001). **(B)** Fold increase in the relative indel ratio (%) between only Cpf1 treated, Cpf1 and dCas9 co-treated, and Cpf1 and nCas9 cotreated samples. Fold change is shown in light gray (dCas9 combination with Cpf1 / only Cpf1 treated), dark gray (nCas9 combination with Cpf1 / only Cpf1 treated) and black (nCas9 combination with Cpf1 / dCas9 combination with Cpf1). **(C)** Relative indel ratio (%) for *DNMT1* plasmid (on-, off-target1, 2) targeted editing with chimeric (cr)RNA guided AsCpf1 in HEK293FT cell. The RNA portion of the Cpf1 (cr)RNA is shown in blue, and the substituted DNA portion is shown in red. X axis indicates various (cr)RNAs for AsCpf1 and Y axis indicates the relative indel ratio (%). Indel ratio (%) is calculated by targeted amplicon sequencing from *DNMT1* site in plasmid (indel frequency (%) = mutant DNA read number / total DNA read number) and normalized to wild-type (cr)RNA (crRNA 1 for *DNMT1*). Data are shown as means ± s.e.m. from three independent experiments. *P*-values are calculated using a twotailed Student’s t-test (ns: not significant, P*:<0.05, P**:<0.01, P***:<0.001, P****:<0.0001). **(D)** Determination of target specificity (on/off cleavage ratio) calculated from (C). Target specificity is shown in black (on/off 1) and gray (on/off 2), respectively.

**Figure 8.**
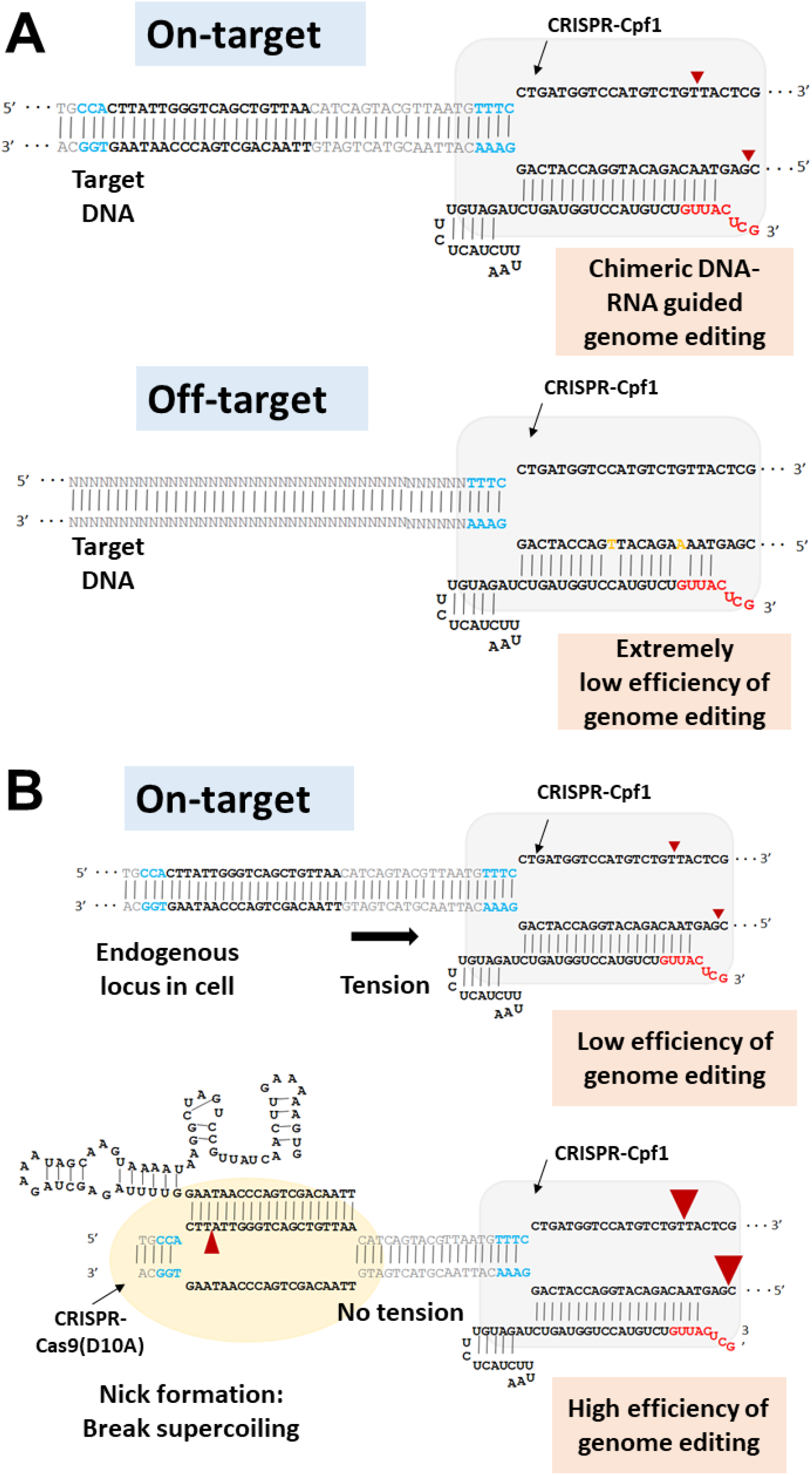
Mechanism underlying the enhanced Cpf1 editing efficiency and specificity of the combination of chimeric DNA-RNA guide and SpCas9 nickase. **(A)** Schematics of the chimeric DNA-RNA-guided editing with Cpf1. Off-target DNA cleavage is dramatically decreased by chimeric (cr)RNA guided Cpf1 binding because of the higher mismatch sensitivity. The red arrowhead indicates the cleavage site for double-strand breaks. **(B)** Enhanced genome editing with a combination of SpCas9 nickase (D10A) and chimeric DNA-RNA guided Cpf1. DNA supercoiling generates a tension that inhibits the chimeric (cr)RNA guided Cpf1 cleavage activity. Nickase relaxed supercoiling enhances chimeric (cr)RNA guided Cpf1 DNA cleavage activity on genomic DNA. The small and large red arrowhead indicates the cleavage site for nick by SpCas9 nickase (D10A) and enhanced double-strand break by Cpf1. Red color in the (cr)RNA indicates DNA substitution. PAM sequences (NGG for SpCas9 nickase and TTTN for AsCpf1) are shown in blue color and mismatched sequence to target sequence are shown in orange color, respectively.

## Discussion

Based on the well-defined interaction of gRNAs and specific amino acids of proteins in the well-established CRISPR-Cas12a complex with target DNA, we screened chimeric guides with high target specificity by reducing off-target cleavage without reducing on-target cleavage efficiency from various (cr)RNAs with DNA substitution at the 5’-end, 3’-end, or seed region. When various chimeric guides and Cpf1 proteins were applied to off-targets sequences which is similar to on-targets, the tendency shows that if mismatch exists in the PAM-proximal seed region, as in the result of the existing off-target detection paper [19, 20], off-target cleavage was hardly generated. But, off-target cleavage with mismatch in the PAM-distal region or middle of the target sequence was observed in the *DNMT1, GRIN2B*, and *EMX1* gene targets. At this time, in the case of continuous (+8nt) DNA substitution at the 3′-end of the guide, off-target cleavage was reduced in *DNMT1* off-target 1, 2 site and *GRIN2B* off-target 2 site, which greatly increases the target specificity of Cpf1. On the other hand, off-target cleavage was not detected or changed, but on-target cleavage efficiency was rather increased for *FANCF* and *EMX1* gene targets. This is thought to be a complex effect of Cpf1’s structural recognition of four non-base pairing regions of (cr)RNA and change of the binding energy in the 20 base paired regions of the target sequence. From a thermodynamic point of view, it can be seen that less off-target cleavage occurs when the hybridization binding energy is unstable, depending on the position of mismatch in the off-target recognized by Cpf1. Unlike the 3’-end, the DNA substitution on the 5’-hairpin seems to have a significant effect on the Cpf1 activity by structural changes in the hairpin region. In addition to the 5’-end hairpin structure, DNA substitution in the seed region inside the target sequence of the RNA guide significantly reduces on-target cleavage activity which indicates that Cpf1 protein requires 2’-OH recognition to stably interact with target DNA - (cr)RNA duplex structure. When the genome editing was induced in the cell using highly accurate chimeric (cr)RNA with 3’-end DNA substitution, we observed a decrease in endonuclease activity which is different from the result of naked PCR amplicon or plasmid cleavage. The recovery of Cpf1 activity suggests that low activity with chimeric DNA-RNA guided Cpf1 is due to the effects of various factors such as exonucleases present in the cell or supercoiled conformation of the intracellular genomic DNA. In this study, chemical modification of the 3’-end of the chimeric guide solved stability issue, and the combination of SpCas9 nickase and Cpf1 solved the supercoiling issue of the chromosome, thus improving target-specific genome editing efficiency. In particular, we can increase the editing efficiency of Cpf1 by using SpCas9 nickase as in our model **(Figure 8)**, while the target specificity is greatly improved because chimeric guides that reduce the hybridization energy and increase the sensitivity to mismatch. And target DNA specificity is further enhanced along with the use of SpCas9 nickase which cannot bind to other off-target sites. By using SpCas9 nickase and chimeric (cr)RNA guided Cpf1 in combination, we have developed an extremely safe genome editing technique that reduces off-target cleavage. Overall, future structural and biochemical studies on chimeric DNA-RNA recognition of the CRISPR-Cas12a protein are required to improve the target DNA cleavage efficiency by protein engineering. In particular, if the target-specific genome editing efficiency can be improved by changing the amino acid residue(s) in Cpf1 required for full DNA recognition, non-specific cleavage can be dramatically reduced *in vivo*, and thus, the safety of target-specific genome editing would increase.

### Conclusions

In this study, we suggested that the target DNA sequence specificity of Cpf1 was improved by partial substitution of the gRNA with DNA. We further increased the possibility of target specific genome editing by enhancing the intracellular operating efficiency of chimeric guide based Cpf1 using the combination of nickase Cas9. The results of this study will aid in the development of chimeric DNA-RNA-guided target-specific Cpf1 that can be applied in various living organisms, including microorganisms, plants, animals, and ultimately, human beings. Obviously, safety issues remain to be addressed in clinical trials.

## Methods

### Preparation of the CRISPR-Cas12a recombinant protein and chimeric guides

For purification of Cpf1 recombinant protein, pET28a-Cpf1 (*Acidaminococcus sp*. (As)Cpf1, *Lachnospiraceae bacterium* (Lb)Cpf1, *Francisella novicida* (Fn)Cpf1) bacterial expression vectors were introduced into E. coli BL21 (DE3) species and transformed, and then cultured at 37°C until O.D. = 0.6. After 48 hours of IPTG inoculation, bacterial cells were precipitated to remove the culture medium, and the remaining precipitate was resuspended with bufferA [20 mM Tris-HCl (pH8.0), 300 mM NaCl, 10 mM β-mercaptoethanol, 1% TritonX-100, 1mM PMSF]. Then, bacterial cell membranes were broken through sonication (ice, 3 min), and cell lysate was harvested after centrifugation (5000 rpm, 10 min). Ni-NTA resin pre-washed with buffer [20 mM Tris-HCl (pH8.0), 300 nM NaCl] and the ultrasonically disrupted intracellular solution were mixed and stirred for 1 hour 30 minutes in cold room (4°C). After bacterial cell precipitation, non-specific binding components were removed by washing with bufferB [20mM Tris-HCl (pH8.0), 300nM NaCl] at 10 times volume, bufferC [20mM Tris-HCl (pH8.0), 300 nM NaCl, 200 mM Imidazole] is used to elute the AsCpf1 protein. The bufferC which is used for protein elution is exchanged to bufferE [200 mM NaCl, 50 mM HEPES (pH 7.5), 1mM DTT, 40% glycerol] by using centricon (Amicon Ultra,) and stored at −80°C. Chimeric DNA-RNA guides (bioneer) were batch synthesized according to the target sequences in each target gene **(Table S1)**.

### In-vitro transcription and purification of the (cr)RNA for Cpf1

For in-vitro transcription, each sense and anti-sense DNA oligo containing the target (cr)RNA sequence **(Table S1)** was purchased (Macrogen). Mix the annealed DNA template with T7 RNA polymerase (NEB) and mixture (50mM MgCl_2_, 100mM rNTP (Jena Bio, NU-1014), 10X RNA polymerase reaction buffer, RNase Inhibitor Murine, 100mM DTT, DEPC), then (cr)RNA is synthesized by incubation for 8 hours at 37°C. After synthesis, the DNA template was completely removed by incubation at 37°C for 1 hour with DNase, and only RNA was separated through the column (MP Biomedicals, GENECLEAN^®^ Turbo Kit). Purified RNA is concentrated through lyophilization (2000 rpm, 1hour, −55°C, 25°C).

### In-vitro DNA cleavage assay and image based calculation of the DNA cleavage efficiency

PCR amplicons are obtained from purified genomic DNA (HEK293FT) using DNA primers **(Table S2)** corresponding to each target locus (*DNMT1, CCR5, FANCF, GRIN2B, EMX1*). To cleavage the amplicons, purified recombinant Cpf1 protein and synthesized chimeric (cr)RNA-DNA (purchased from Bioneer) or purified (cr)RNA corresponding to each locus are premixed and incubated at 37°C for 1 hour in cleavage buffer (NEB3, 10ul volumn). Then, stop the reaction by adding a stop buffer (100mM EDTA, 1.2% SDS), and check the DNA cleavage by 2% agarose gel electrophoresis. DNA cleavage efficiency was determined by calculating the image pattern according to the formula (Intensity of the cleaved fragment / total sum of the fragment intensity X 100 =%) measured using imageJ software(NIH).

### Cell sub-culture and transfection

HEK293FT cell line (ATCC) was passaged in DMEM media (DMEM (Gibco) with 10% FBS (Gibco)) every 48 hours at 37°C, 5% CO2 to maintain a confluency of 70%. For cell transfection, electroporation kit (Lonza, V4XC-2032) was used, and 10^5 cells were mixed with Cpf1-chimeric guide pre-mixed complex(Cpf1: 60 pmol, (cr)RNA: 240 pmol) and followed by electric shock (program: CM-130) in electroporation buffer(manufactor’s guide). Plasmids (*DNMT1*, *DNMT1-off1, off2*, 1μg) were cotransfected with Cpf1-chimeric guide pre-mixture for related experiments. Subsequently, transfer the transfected cells into DMEM media solution (500ul) of a 24-well plate pre-incubated for 30 minutes at 37°C and 5% CO2, and incubate at the same conditions (37°C and 5% CO2).

### Genomic DNA purification and T7E1 assay

After delivery of the chimeric (cr)RNA-DNA and the recombinant Cpf1 protein complex into cells (HEK293FT), genomic DNA is isolated using a genomic DNA purification kit (Qiagen, DNeasy Blood & Tissue Kit) at 48hours later. From this, PCR amplicons (*DNMT1, CCR5, FANCF, GRIN2B, EMX1*) are obtained using DNA primers **(Table S2)** corresponding to each locus, and denature-reannealing (manufactor’s protocol) is performed with a PCR machine. Amplicon is incubated at 37°C for 25min in cleavage buffer (NEBuffer 2, 10ul volumn) using purified recombinant T7E1 enzyme (NEB, T7 endonuclease I). After T7E1 reaction, the reaction was stopped by adding a stop buffer (100 mM EDTA, 1.2% SDS), and DNA cleavage was confirmed by 2% agarose gel electrophoresis.

### Targeted deep sequencing and data analysis

Obtain PCR amplicons (*DNMT1, CCR5, FANCF, GRIN2B, EMX1*) using DNA primer **(Table S2)** corresponding to the target locus. Then, nested PCR (Denaturation: 98°C – 30 sec, Primer annealing: 58°C – 30 sec, Elongation: 72°C – 30 sec, 35 cycles) was performed to insert the adapter and index sequences into both 5’ and 3’-end of the amplicon (Denaturation: 98°C – 30 sec, Primer annealing: 62°C – 15 sec, Elongation: 72°C – 15 sec, 35 cycles). Thereafter, the tagged amplicon mixture was loaded onto a mini-SEQ analyzer(illumina, SY-420-1001) according to the manufactor’s guidelines and subjected to targeted deep sequencing. The saved Fastq files were analyzed with Cas-Analyzer[30] and the indel efficiency was calculated.

### GUIDE-seq and data analysis

Guide-seq was performed as shown in a previous study[25]. In short, HEK293T cells were nucleofected using electroporation kit (Lonza, V4XC-2032) following manufacturer’s guide. 100 pmol of double stranded oligonucleotides (dsODNs) (100 pmol) were co-nucleofected with Cpf1-guide pre-mixed complex in a 24-well plate at 10^5 cells per well. Genomic DNA was extracted using a genomic DNA purification kit (Qiagen, DNeasy Blood & Tissue Kit) at 48hours after nucleofection. The isolated genomic DNA was fragmented via sonication (Covaris) to around 500bp and was purified with AMPure XP beads (Beckman Coulter). Next, the fragmented DNA was treated with NEBNext End Prep (NEB) and then was subjected to adaptor ligation for Illumina using NEBNext Adaptor for Illumina (NEB). Then USER Enzyme (NEB) was added the DNA and the treated DNA was purified using AMPure XP beads. In order to obtain the sequence information of dsODN insertion sites, 2 rounds of PCR was conducted using DNA primer corresponding to the adaptor sequence and the both 5’ and 3’-ends of dsODN sequence. Then, another nested PCR was performed to add the index sequences for deep sequencing by Illumina platform. The deep sequencing results were analyzed using the publicly available python-based analysis pipeline. The reads were first mapped to the reference genome and then the information on off-target sites was derived based on the sequence of the guide RNA.

## Ethics approval and consent to participate

Not applicable.

## Consent for publication

Not applicable.

## Competing interests

The authors declare that they have no competing interests.

## Funding

This research was supported by grants from the National Research Foundation (NRF) funded by the Korean Ministry of Education, Science and Technology (NRF-2019R1C1C1006603). The study was also supported by the grants from the Korea Research Institute of Bioscience & Biotechnology (KRIBB; Research Initiative Program KGM4251824, KGM5381911 and KGM1051911).

## Authors’ contributions

Conceptualization, H.K., W.L. and S.H.L.; Methodology, J.K.H. and S.H.L.; Software, H.K., W.L., J.K.H. and S.H.L.; Validation, H.K. and W.L.; Formal Analysis, H.K. and W.L.; Investigation, H.K., W.L., and S.H.K., H.L., W.J.S.; Resources, K.S.L., Y.H.P., B.S.S., and Y.B.J; Data Curation, H.K., W.L., H.L., W.J.S. and S.H.L.; Writing-Original Draft, H.K., W.L. and S.H.L.; Writing-Review & Editing, B.H.J., D.S.L., S.U.K. and S.H.L.; Visualization, H.K., W.L., Y.H.P and S.H.L.; Supervision, S.U.K. and S.H.L.; Project Administration, D.S.L., S.U.K. and S.H.L.; Funding Acquisition, S.U.K. and S.H.L.

## Acknowledgements

Authors thank to the other members of the National Primate Research Center (NPRC) for their helpful discussions.

