## Supplementary Information for "Enhancement of Target Specificity of CRISPR-Cas12a by Using a Chimeric DNA-RNA Guide"

### Supplementary Tables

#### Table S1. Sequence information for chimeric DNA-RNA guides used in this study.

| Target gene (guide no.) | CRISPR-Cas12a (AsCpf1) target sequence (5'-3') | crRNA sequence for AsCpf1 (5'-3') |
|---|---|---|
| hDNMT1 **crRNA1** | TTTCCTGATGGTCCATGTCTGTTACTCG | **5'AAUUUCUACUCUUGUAGAUCUGAUGGUCCAUGUCUGUUACUCG** 3' |
| hDNMT1 **crRNA2** | TTTCCTGATGGTCCATGTCTGTTACTCG | **5'AAUUUCUACUCUUGUAGAUCUGAUGGUCCAUGUCUGUUACTCG 3'** |
| hDNMT1 **crRNA3** | TTTCCTGATGGTCCATGTCTGTTACTCG | **5'AAUUUCUACUCUUGUAGAUCUGAUGGUCCAUGUCUGTTACTCG 3'** |
| hDNMT1 **crRNA3-2** | TTTCCTGATGGTCCATGTCTGTTACTCG | **5'AAUUUCUACUCUUGUAGAUCUGAUGGUCCAUGUCTGTTACTCG 3'** |
| hDNMT1 **crRNA3-3** | TTTCCTGATGGTCCATGTCTGTTACTCG | **5'AAUUUCUACUCUUGUAGAUCUGAUGGUCCAUGUCTGTTACTCG 3'** |
| hDNMT1 **crRNA3-4** | TTTCCTGATGGTCCATGTCTGTTACTCG | **5'AAUUUCUACUCUUGUAGAUCUGAUGGUCCAUGTCTGTTACTCG 3'** |
| hDNMT1 **crRNA4** | TTTCCTGATGGTCCATGTCTGTTACTCG | **5'AAUUUCUACUCUUGUAGAUCUGAUGGUCCAUGTCTGTTACTCG 3'** |
| hDNMT1 **crRNA5** | TTTCCTGATGGTCCATGTCTGTTACTCG | **5'AAUUUCUACUCUUGUAGAUCUGAUGGUCCATGTCTGTTACTCG 3'** |
| hDNMT1 **crRNA6** | TTTCCTGATGGTCCATGTCTGTTACTCG | **5'AAUUUCUACUCUUGUAGAUCUGATGGTCCATGTCTGTTACTCG 3'** |
| hDNMT1 **crRNA7** | TTTCCTGATGGTCCATGTCTGTTACTCG | **5'AAUUUCUACUCUUGUAGAUCTGATGGTCCATGTCTGTTACTCG 3'** |
| hDNMT1 **crRNA8** | TTTCCTGATGGTCCATGTCTGTTACTCG | **5'AATTTCTACTCTTGTAGATCTGATGGTCCATGTCTGTTACTCG 3'** |
| hDNMT1 **crRNA9** | TTTCCTGATGGTCCATGTCTGTTACTCG | **5'AATTUCUACUCUUGUAGAUCUGAUGGUCCAUGUCUGUUACUCG 3'** |
| hDNMT1 **crRNA10** | TTTCCTGATGGTCCATGTCTGTTACTCG | **5'AATTTCTACUCUUGUAGAUCUGAUGGUCCAUGUCUGUUACUCG 3'** |
| hDNMT1 **crRNA11** | TTTCCTGATGGTCCATGTCTGTTACTCG | **5'AATTTCTACTCTUGUAGAUCUGAUGGUCCAUGUCUGUUACUCG 3'** |
| hDNMT1 **crRNA12** | TTTCCTGATGGTCCATGTCTGTTACTCG | **5'AATTTCTACTCTTGTAGAUCUGAUGGUCCAUGUCUGUUACUCG 3'** |
| hDNMT1 | TTTCCTGATGGTCCATGTCTGTTAC | **5'AATTTCTACTCTTGTAGATCUGAUGGUCCAUGUCUGUUA** |

| | | |
|---|---|---|
| **crRNA13** | TCG | **CUCG 3'** |
| hDNMT1 **crRNA14** | TTTCCTGATGGTCCATGTCTGTTACTCG | **5'AAUUUCUACUCUUGUAGAUCTGAUGGUCCAUGUCUGUUACUCG 3'** |
| hDNMT1 **crRNA15** | TTTCCTGATGGTCCATGTCTGTTACTCG | **5'AAUUUCUACUCUUGUAGAUCTGATGGTCCAUGUCUGUUACUCG 3'** |
| hDNMT1 **crRNA16** | TTTCCTGATGGTCCATGTCTGTTACTCG | **5'AAUUUCUACUCUUGUAGAUCTGATGGTCCATGUCUGUUACUCG 3'** |
| hDNMT1 **crRNA17** | TTTCCTGATGGTCCATGTCTGTTACTCG | **5'AAUUUCUACUCUUGUAGAUCUGAUGGUCCAUGUCUGUUACUCG 3'** |
| hDNMT1 **crRNA18** | TTTCCTGATGGTCCATGTCTGTTACTCG | **5'AAUUUCUACUCUUGUAGAUCTGAUGGUCCAUGUCUGUUACUCG 3'** |
| hDNMT1 **crRNA19** | TTTCCTGATGGTCCATGTCTGTTACTCG | **5'AAUUUCUACUCUUGUAGAUCUGAUGGUCCAUGUCUGUUACUCG 3'** |
| hDNMT1 **crRNA20** | TTTCCTGATGGTCCATGTCTGTTACTCG | **5'AAUUUCUACUCUUGUAGAUCUGAUGGUCCAUGUCUGUUACUCG 3'** |
| hDNMT1 **crRNA21** | TTTCCTGATGGTCCATGTCTGTTACTCG | **5'AAUUUCUACUCUUGUAGAUCUGATGGUCCAUGUCUGUUACUCG 3'** |
| hDNMT1 **crRNA22** | TTTCCTGATGGTCCATGTCTGTTACTCG | **5'AAUUUCUACUCUUGUAGAUCUGAUGGUCCAUGUCUGUUACUCG 3'** |
| hDNMT1 **crRNA23** | TTTCCTGATGGTCCATGTCTGTTACTCG | **5'AAUUUCUACUCUUGUAGAUCUGAUGGUCCAUGUCUGUUACUCG 3'** |
| hDNMT1 **crRNA24** | TTTCCTGATGGTCCATGTCTGTTACTCG | **5'AAUUUCUACUCUUGUAGAUCUGAUGGTCCAUGUCUGUUACUCG 3'** |
| hDNMT1 **crRNA25** | TTTCCTGATGGTCCATGTCTGTTACTCG | **5'AAUUUCUACUCUUGUAGAUCUGAUGGUCCAUGUCUGUUACUCG 3'** |
| hDNMT1 **crRNA26** | TTTCCTGATGGTCCATGTCTGTTACTCG | **5'AAUUUCUACUCUUGUAGAUCUGAUGGUCCAUGUCUGUUACUCG 3'** |
| hDNMT1 **crRNA27** | TTTCCTGATGGTCCATGTCTGTTACTCG | **5'AAUUUCUACUCUUGUAGAUCUGAUGGUCCAUGUCUGUUACUCG 3'** |
| hCCR5 **crRNA51** | TTTGTGCACAGGGTGGAACAAGATGGAT | **5'AAUUUCUACUCUUGUAGAUUGCACAGGGUGGAACAAGAUGGAU 3'** |
| hCCR5 **crRNA52** | TTTGTGCACAGGGTGGAACAAGATGGAT | **5'AAUUUCUACUCUUGUAGAUUGCACAGGGUGGAACAAGAUGGAT 3'** |
| hCCR5 **crRNA53** | TTTGTGCACAGGGTGGAACAAGATGGAT | **5'AAUUUCUACUCUUGUAGAUUGCACAGGGUGGAACAAGATGGAT 3'** |
| hCCR5 **crRNA54** | TTTGTGCACAGGGTGGAACAAGATGGAT | **5'AAUUUCUACUCUUGUAGAUUGCACAGGGUGGAACAAGATGGAT 3'** |
| hCCR5 **crRNA55** | TTTGTGCACAGGGTGGAACAAGATGGAT | **5'AAUUUCUACUCUUGUAGAUUGCACAGGGTGGAACAAGATGGAT 3'** |
| hCCR5 **crRNA56** | TTTGTGCACAGGGTGGAACAAGATGGAT | **5'AAUUUCUACUCUUGUAGAUUGCACAGGGTGGAACAAGATGGAT 3'** |
| hCCR5 **crRNA57** | TTTGTGCACAGGGTGGAACAAGATGGAT | **5'AAUUUCUACUCUUGUAGAUTGCACAGGGTGGAACAAGATGGAT 3'** |
| hCCR5 **crRNA58** | TTTGTGCACAGGGTGGAACAAGATGGAT | **5'AATTTCTACTCTTGTAGATTGCACAGGGTGGAACAAGATGGAT 3'** |
| hCCR5 **crRNA61** | TTTGTGCACAGGGTGGAACAAGATGGAT | **5'AAUUUCUACUCUUGUAGAUTGCACAGGGUGGAACAAGAUGGAU 3'** |
| hCCR5 **crRNA62** | TTTGTGCACAGGGTGGAACAAGATGGAT | **5'AAUUUCUACUCUUGUAGAUUGCACAGGGUGGAACAAGAUGGAU 3'** |
| hCCR5 **crRNA63** | TTTGTGCACAGGGTGGAACAAGATGGAT | **5'AAUUUCUACUCUUGUAGAUUGCACAGGGUGGAACAAGAUGGAU 3'** |
| hCCR5 **crRNA64** | TTTGTGCACAGGGTGGAACAAGATGGAT | **5'AAUUUCUACUCUUGUAGAUUGCACAGGGUGGAACAAGAUGGAU 3'** |
| hCCR5 **crRNA65** | TTTGTGCACAGGGTGGAACAAGATGGAT | **5'AAUUUCUACUCUUGUAGAUUGCACAGGGUGGAACAAGAUGGAU 3'** |
| hCCR5 | TTTGTGCACAGGGTGGAACAAGAT | **5'AAUUUCUACUCUUGUAGAUUGCACAGGGUGGAACAAG** |

| | | |
|---|---|---|
| **crRNA66** | GGAT | **AUGGAU 3'** |
| hCCR5 **crRNA67** | TTTGTGCACAGGGTGGAACAAGAT GGAT | **5'AAUUUCUACUCUUGUAGAUUGCACAGGGUGGAACAAG AUGGAU 3'** |
| hCCR5 **crRNA68** | TTTGTGCACAGGGTGGAACAAGAT GGAT | **5'AAUUUCUACUCUUGUAGAUUGCACAGGGUGGAACAAG AUGGAU 3'** |
| hCCR5 **crRNA69** | TTTGTGCACAGGGTGGAACAAGAT GGAT | **5'AAUUUCUACUCUUGUAGAUUGCACAGGGUGGAACAAG AUGGAU 3'** |
| hCCR5 **crRNA70** | TTTGTGCACAGGGTGGAACAAGAT GGAT | **5'AAUUUCUACUCUUGUAGAUUGCACAGGGTGGAACAAGA UGGAU 3'** |
| hFANCF **crRNA81** | TTTGGTCGGCATGGCCCCATTCGC ACGG | **5'AAUUUCUACUCUUGUAGAUGUCGGCAUGGCCCCAUUCG CACGG 3'** |
| hFANCF **crRNA82** | TTTGGTCGGCATGGCCCCATTCGC ACGG | **5'AAUUUCUACUCUUGUAGAUGUCGGCAUGGCCCCAUUCG CACGG 3'** |
| hFANCF **crRNA82-2** | TTTGGTCGGCATGGCCCCATTCGC ACGG | **5'AAUUUCUACUCUUGUAGAUGUCGGCAUGGCCCCAUTCG CACGG 3'** |
| hGRIN2B **crRNA83** | TTTGGTGCTCAATGAAAGGAGATAA GGT | **5'AAUUUCUACUCUUGUAGAUGUGCUCAAUGAAAGGAGAU AAGGU 3'** |
| hGRIN2B **crRNA84** | TTTGGTGCTCAATGAAAGGAGATAA GGT | **5'AAUUUCUACUCUUGUAGAUGUGCUCAAUGAAAGGAGAU AAGGT 3'** |
| hGRIN2B **crRNA84-2** | TTTGGTGCTCAATGAAAGGAGATAA GGT | **5'AAUUUCUACUCUUGUAGAUGUGCUCAAUGAAAGGAGAT AAGGT 3'** |
| hEMX1 **crRNA85** | TTTGTCCTCCGGTTCTGGAACCACA CCT | **5'AAUUUCUACUCUUGUAGAU**UCCUCCGGUUCUGGAACCA CACCU 3'** |
| hEMX1 **crRNA86** | TTTGTCCTCCGGTTCTGGAACCACA CCT | **5'AAUUUCUACUCUUGUAGAU**UCCUCCGGUUCUGGAACCA CACCT 3'** |
| hEMX1 **crRNA86-2** | TTTGTCCTCCGGTTCTGGAACCACA CCT | **5'AAUUUCUACUCUUGUAGAU**UCCUCCGGUUCUGGAACCA CACCT 3'** |
| hDNMT1 **crRNA2-PS** | TTTCCTGATGGTCCATGTCTGTTAC TCG | **5'AAUUUCUACUCUUGUAGAUCUGAUGGUCCAUGUCUGUU ACTCG (PS)3'** |
| hDNMT1 **crRNA3-PS** | TTTCCTGATGGTCCATGTCTGTTAC TCG | **5'AAUUUCUACUCUUGUAGAUCUGAUGGUCCAUGUCUGTT ACTCG (PS)3'** |

PAM sequences (TTTN) for AsCpf1 in the target DNA are shown in blue and substituted DNA sequences in (cr)RNA targets are shown in red, respectively. PS indicates a 3'-end modification of the (cr)RNA with phosphorothioate.

### Table S2. Sequence information for DNA primers used in this study.

| **Target gene (primer direction)** | **DNA sequence (5' to 3')** |
|---|---|
| hDNMT1 on-target (F1) | GTTGCACGTGTCAAGTGCTTA |
| hDNMT1 on-target R1 | TTAAAATCCAGAATGCACAAAGTACT |
| hDNMT1 on-target F2 | GTGAATTTGGCTCAGCAGGCA |
| hDNMT1 on-target R2 | AAGCGAACCTCACACAACAGC |
| hDNMT1 off-target1 F1 | TTGACGGCAGTATTACAGGTAG |
| hDNMT1 off-target1 R1 | AAGGTCAAATGCCGTTTAACCA |
| hDNMT1 off-target1 F2 | GTAGTCAGGCATGAGTGGCA |
| hDNMT1 off-target1 R2 | TGCCTCTTTCCCAGGATTCT |
| hDNMT1 off-target2 F1 | TCTCAGGCAAGTCACAACTCT |
| hDNMT1 off-target2 R1 | TAGGCACATGAAGGTCAAATGC |
| hDNMT1 off-target2 F2 | GTTACAGGTAGTTAAGCAGGCA |
| hDNMT1 off-target2 R2 | AAATGCCATATTTAACCGTGATCCT |
| hDNMT1 off-target3 F1 | TCATGCCTTTCTGGGTCTCAT |
| hDNMT1 off-target3 R1 | CACCTGACCTCTGTCACTTTA |
| hDNMT1 off-target3 F2 | CTGTTCAGGGAATGGAAAGTGA |
| hDNMT1 off-target3 R2 | CTACATCTACGCTCTCCCCA |

| | |
|---|---|
| hCCR5 on-target F1 | AAGGCTGAGCTGCACCATGC |
| hCCR5 on-target R1 | AGGATGATGAAGAAGATTCCA |
| hCCR5 on-target F2 | CATTCACTCCATGGTGCTATA |
| hCCR5 on-target R2 | ATAGAGCCCTGTCAAGAGTTGA |
| hCCR5 off-target1 F1 | AGAAATAGGAGTCTTCATGCCC |
| hCCR5 off-target1 R1 | TGGGGTCAATGAGAGGAGTATT |
| hCCR5 off-target1 F2 | TAGCAAGGAACCTCAAAGTGC |
| hCCR5 off-target1 R2 | CTGCTTCCAATACAATCCACAC |
| hCCR5 off-target2 F1 | TAGTGCTGAAAGCTCAGAGAG |
| hCCR5 off-target2 R1 | TTCGAGAGCCACACATGAAG |
| hCCR5 off-target2 F2 | AGAGAGGGGGTCATAGACTTT |
| hCCR5 off-target2 R2 | ACCTTCCAGCCGGTATATTAAT |
| hFANCF on-target F1 | TGAAAGCGGAAGTAGGGCCT |
| hFANCF on-target R1 | CTCCGCCTGGGTCTTCATCA |
| hFANCF on-target F2 | GGAAGTAGGGCCTTCGCGCA |
| hFANCF on-target R2 | CCTCCTGGAGATTTGGGTTC |
| hFANCF off-target1 F1 | CCCCGCTGAGATCATACTAT |
| hFANCF off-target1 R1 | GAATGGAAGTTTGAGGGTAG |
| hFANCF off-target1 F2 | GGTCCAGACACTAACAATTC |
| hFANCF off-target1 R2 | GATTGCTCACAAATATGGGA |
| hFANCF off-target2 F1 | TGGGGTAGGTCTTCAGGAAA |
| hFANCF off-target2 R1 | CGTTCTAAATTCTCTACAGTCAAC |
| hFANCF off-target2 F2 | TCGTGCTAAGTTACCGAGTT |
| hFANCF off-target2 R2 | GCAATAGTTTTGAGCCAGTGA |
| hGRIN2B on-target F1 | ACCTCTGCTGAGCACGTTTT |
| hGRIN2B on-target R1 | GACAGCAATGCCAATGCTGG |
| hGRIN2B on-target F2 | CTCACTTTGTCTGGCCTTGC |
| hGRIN2B on-target R2 | CTTCTGAGAACGAGCTCTGC |
| hGRIN2B off-target1 F1 | TGACTGCAATTTTGGGTTCCATC |
| hGR1IN2B off-target1 R1 | GCCACTGCCATTTATTATGTAAC |
| hGRIN2B off-target1 F2 | GTCTAATTACACTTGCCATACCT |
| hGRIN2B off-target1 R2 | GAAATCCAGCTGTGACTAAATATG |
| hGRIN2B off-target2 F1 | GGAGCACAATGAGTGTTTTT |
| hGRIN2B off-target2 R1 | GTCTTTTCCTGGTCACATGG |
| hGRIN2B off-target2 F2 | GCCAGAAATGTTATTCCATAGTAG |
| hGRIN2B off-target2 R2 | CAGTGTTCTTACATATCAGACAC |
| hEMX1 on-target F1 | ACTACTCACATCCACTCTGTGAAG |
| hEMX1 on-target R1 | GAGTGGCCAGAGTCCAGCTT |
| hEMX1 on-target F2 | TAGAGGAGCTAGGATGCACAGCA |
| hEMX1 on-target R2 | GCAGCAAGCAGCACTCTGCC |
| hEMX1 off-target1 F1 | CAGAGCCTGGAGAATTGATAG |
| hEMX1 off-target1 R1 | CTGGGGAGCAGAATAAATTATTG |
| hEMX1 off-target1 F2 | GAATAGACCTGGGATGTGCAG |
| hEMX1 off-target1 R2 | GGAACCTGAAGAATGGGATTG |
| hEMX1 off-target2 F1 | AGATTGAGATCATCCACCCT |
| hEMX1 off-target2 R1 | AAAACAGAGAGGTTGAGTCTC |
| hEMX1 off-target2 F2 | TGTCCACCATGTCCCAGAAG |
| hEMX1 off-target2 R2 | ACAAAGAAGGGCATCTCCAG |
| hDNMT1_Adaptor_F | ACACTCTTTCCCTACACGACGCTCTTCCGATCTAGTGTTCAGTCTCCGTGAACGTTC |
| hDNMT1_Adaptor_R | GTGACTGGAGTTCAGACGTGTGCTCTTCCGATCTTCCTTAGCAGCTTCCTCCTCCTT |
| hCCR5_Adaptor_F | ACACTCTTTCCCTACACGACGCTCTTCCGATCTAACAGTTTGCATTCATGGAGGGC |
| hCCR5_Adaptor_R | GTGACTGGAGTTCAGACGTGTGCTCTTCCGATCTAGTTTATCAGGATGAGGATGAC C |
| hFANCF_Adaptor_F | ACACTCTTTCCCTACACGACGCTCTTCCGATCTCACTACCTACGTCAGCACCTGGG ACC |
| hFANCF_Adaptor_R | GTGACTGGAGTTCAGACGTGTGCTCTTCCGATCTGGAAGTTCGCTAATCCCGGAAC TGGA |
| hGRIN2B_Adaptor_F | ACACTCTTTCCCTACACGACGCTCTTCCGATCTCTCTCATTCTGCAGAGCAAATACC AGAGAT |
| hGRIN2B_Adaptor_R | GTGACTGGAGTTCAGACGTGTGCTCTTCCGATCTCCTGCAAACACAAAGAAAGAGC ATGTTAAA |

| | |
|---|---|
| hEMX1_Adaptor_F | ACACTCTTTCCCTACACGACGCTCTTCCGATCTGGCCTCCTGAGTTTCTCATCTGTGC |
| hEMX1_Adaptor_R | GTGACTGGAGTTCAGACGTGTGCTCTTCCGATCTCCTGCTTCGTGGCAATGCGCCAC |

Sequences of the forward and reverse adapter primers used in next-generation sequencing are shown in green and blue, respectively.

**Table S3. Sequence information for the sgRNA for dead and nickase (D10A) SpCas9 used in this study.**

| **Target gene (guide no.)** | **CRISPR-Cas9 target sequence (5'-3')** | **sgRNA sequence for dead or nickase SpCas9 (5'-3')** |
|---|---|---|
| hDNMT1 **sgRNA1** | **TTAACAGCTGACCCAATAAGTGG** | **5'GTTAACAGCTGACCCAATAAGGTTTTAGAGCTAGAAATAGCAAGTTAAAATAAGGCTAGTCCGTTATCAACTTGAAAAAGTGGCACCGAGTCGGTGC**3' |

PAM sequence (NGG) in the target DNA for the SpCas9 effector is shown in blue.

### Supplementary Figures

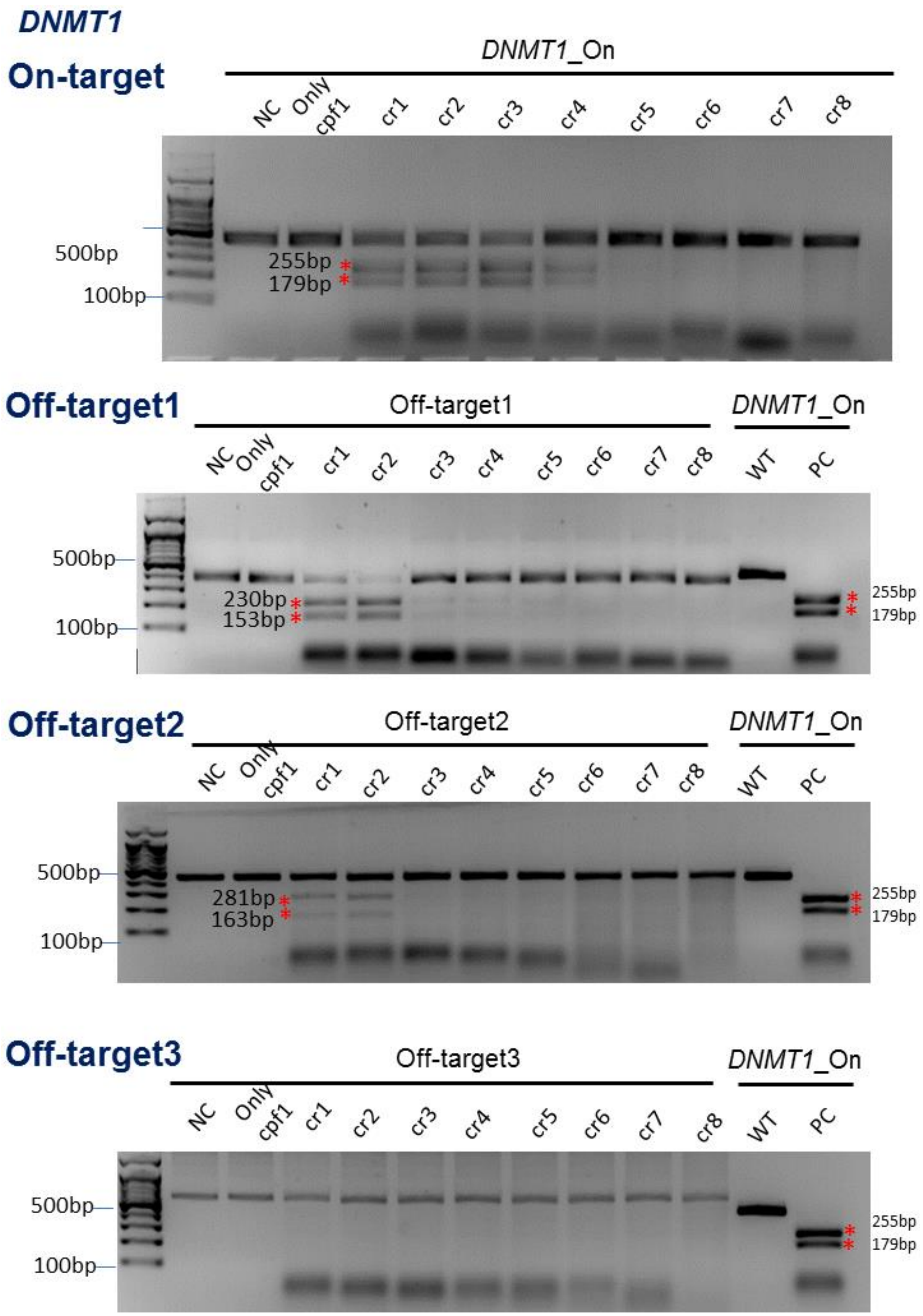

### *DNMT1*

#### On-target

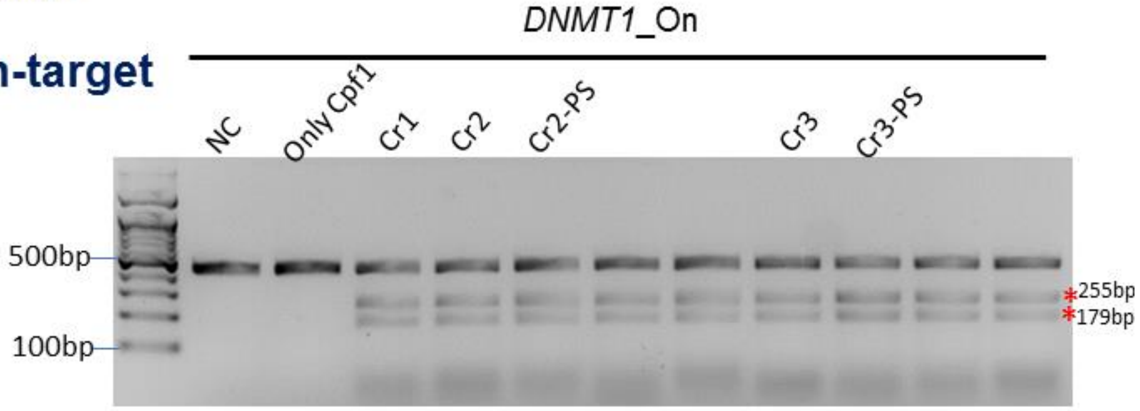


#### Off-target1

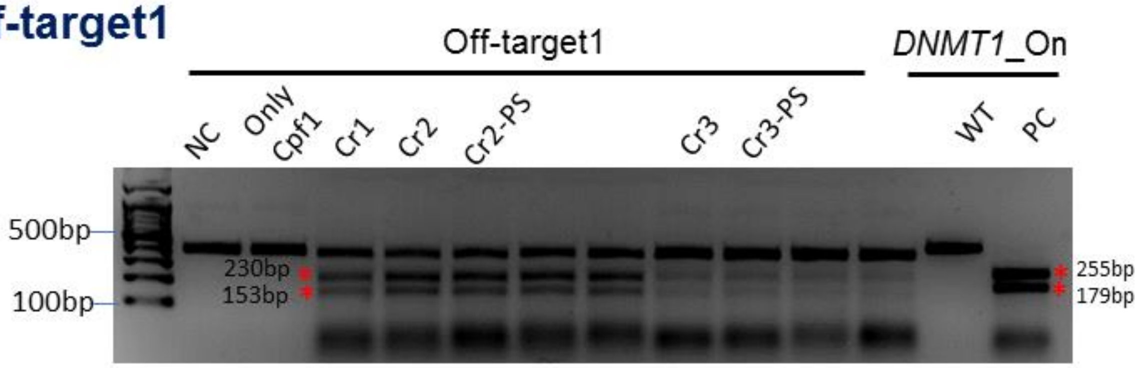


#### Off-target2

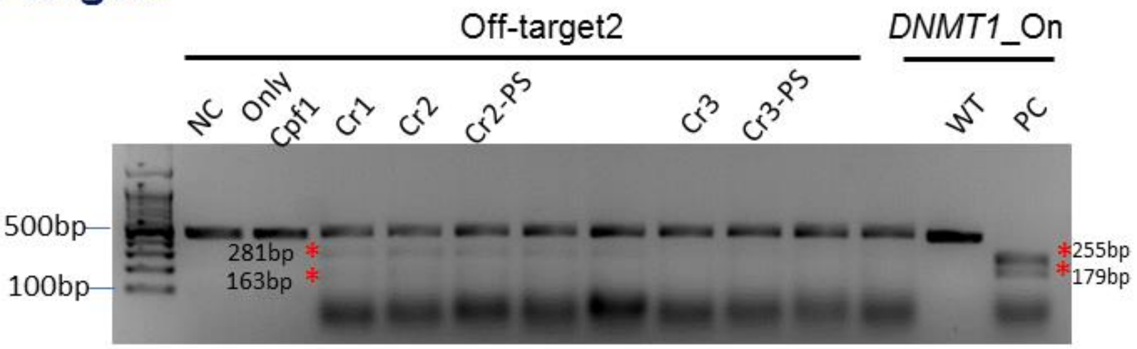


#### Off-target3

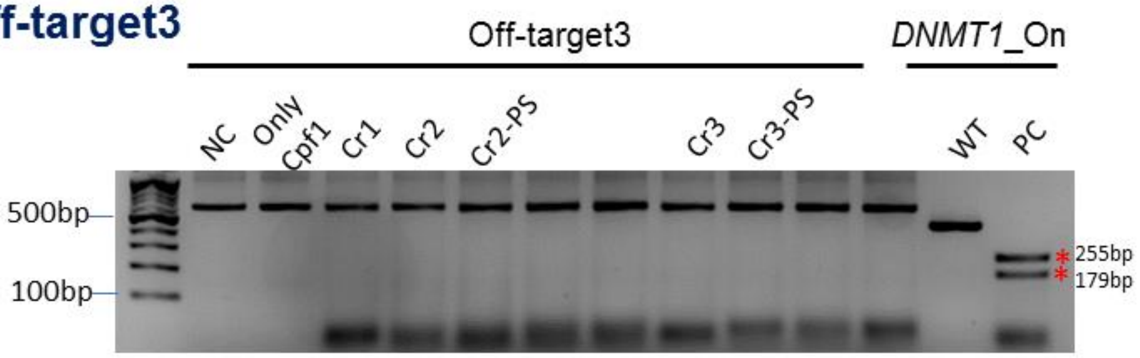

***DNMT1***

**On-target**

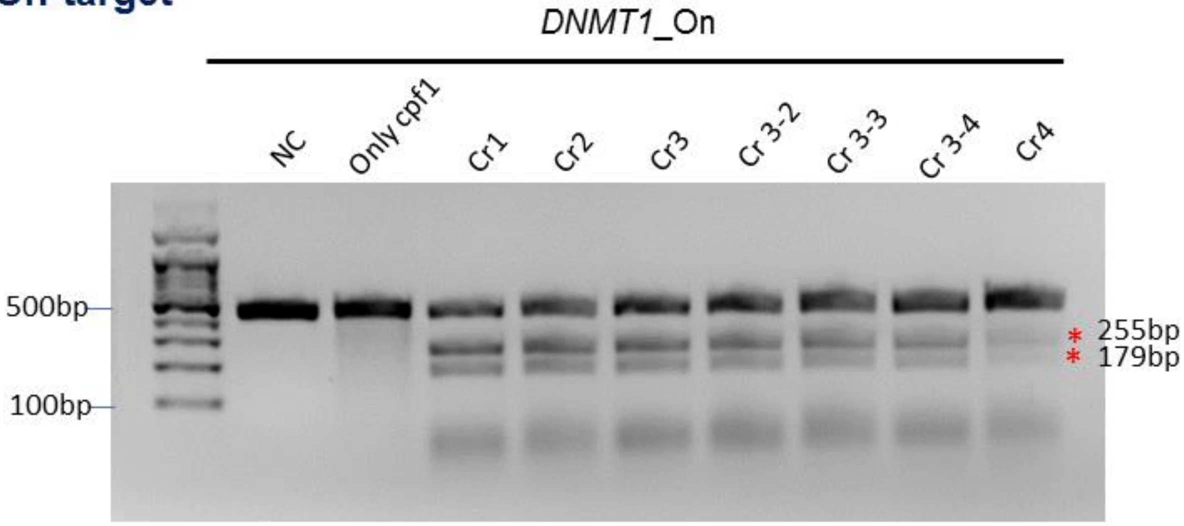


***DNMT1***

**On-target**

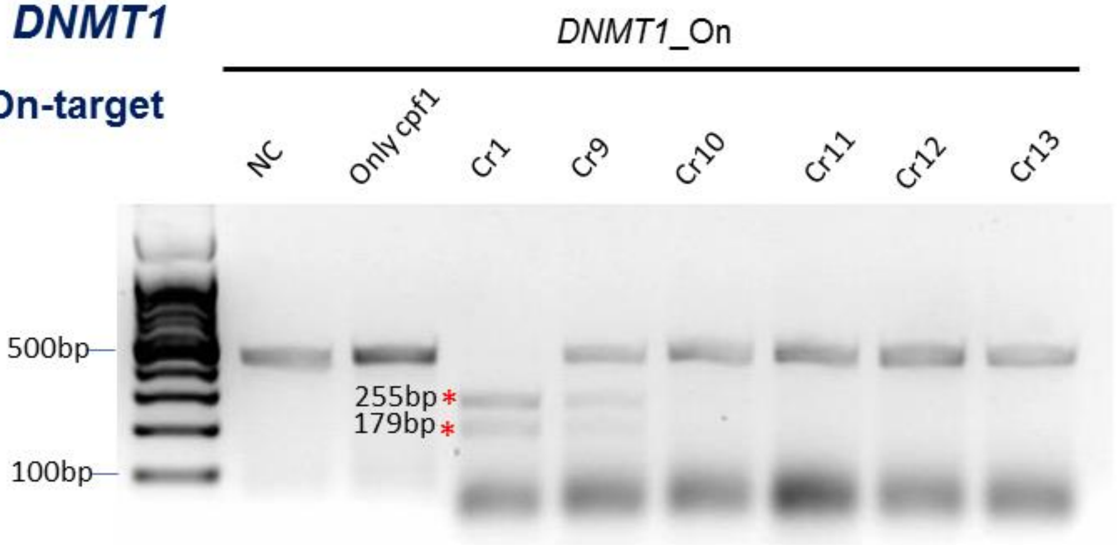

### *DNMT1*

### On-target

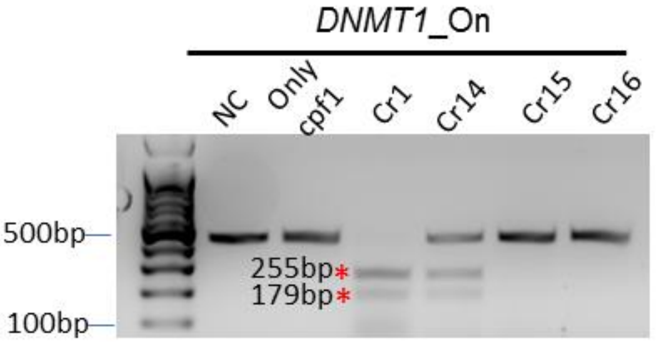


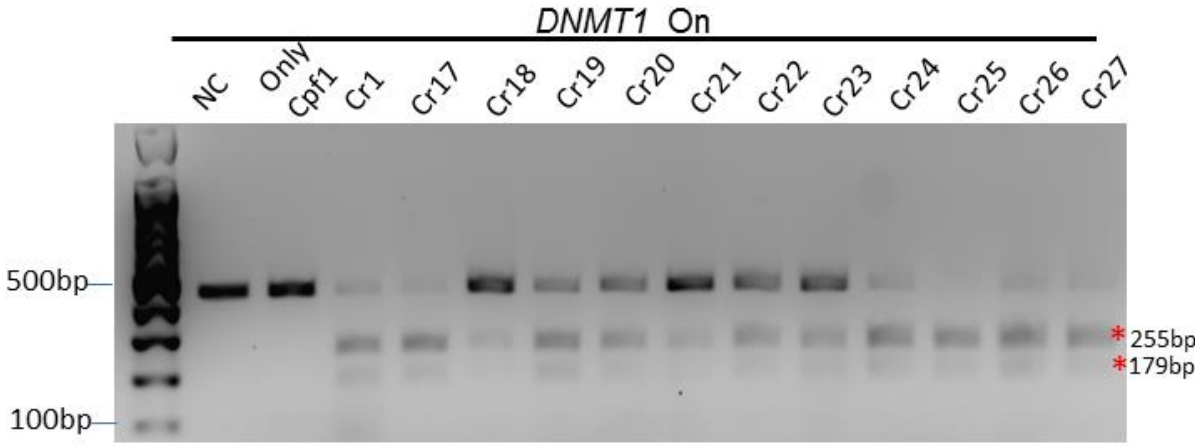


### Off-target1

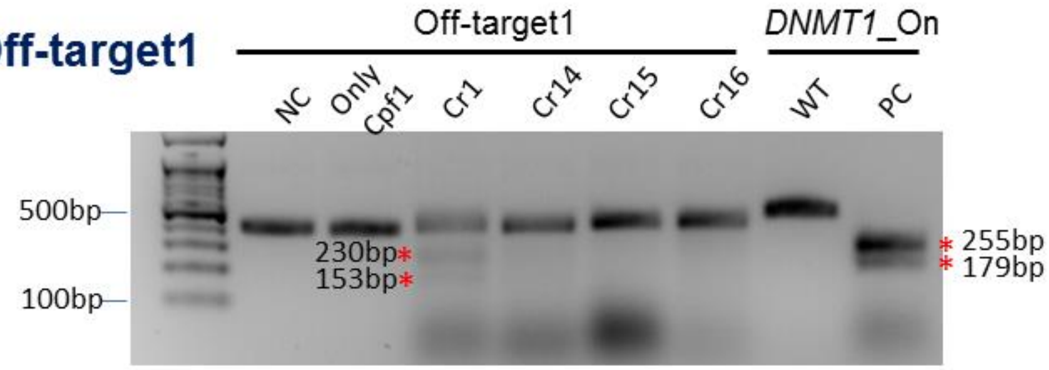


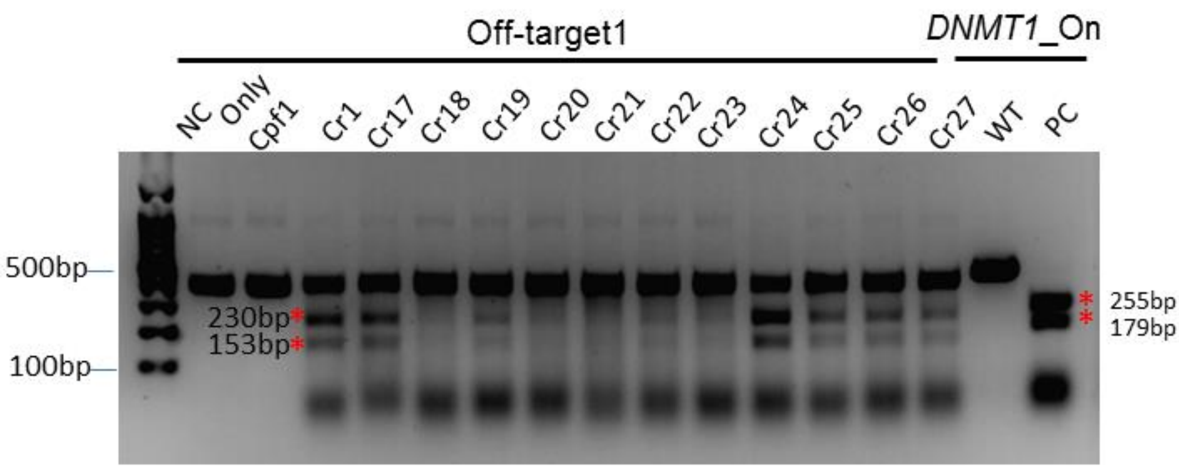

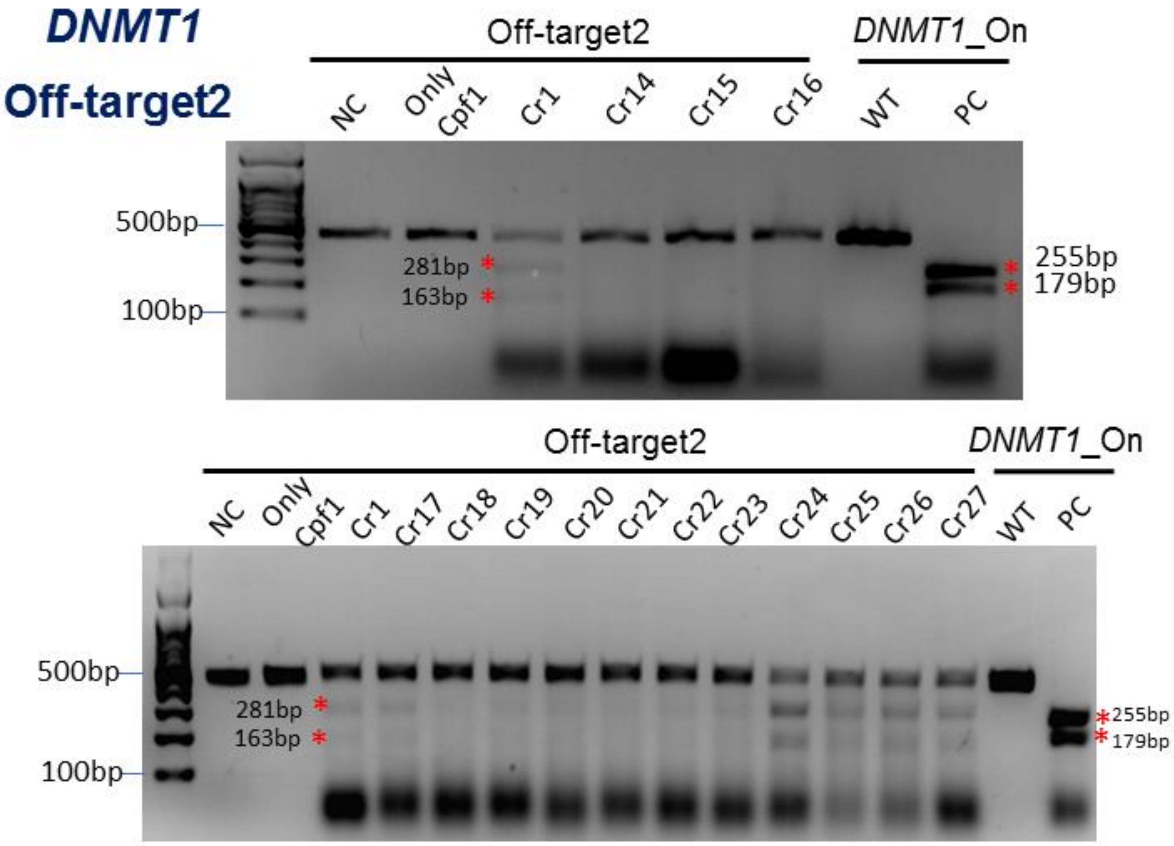
DNMT1
Off-target2
Off-target2
DNMT1_On
NC
Only Cpf1
Cr1
Cr14
Cr15
Cr16
WT
PC
500bp
100bp
281bp
163bp
255bp
179bp
Off-target2
DNMT1_On
NC
Only Cpf1
Cr1
Cr17
Cr18
Cr19
Cr20
Cr21
Cr22
Cr23
Cr24
Cr25
Cr26
Cr27
WT
PC
500bp
100bp
281bp
163bp
255bp
179bp

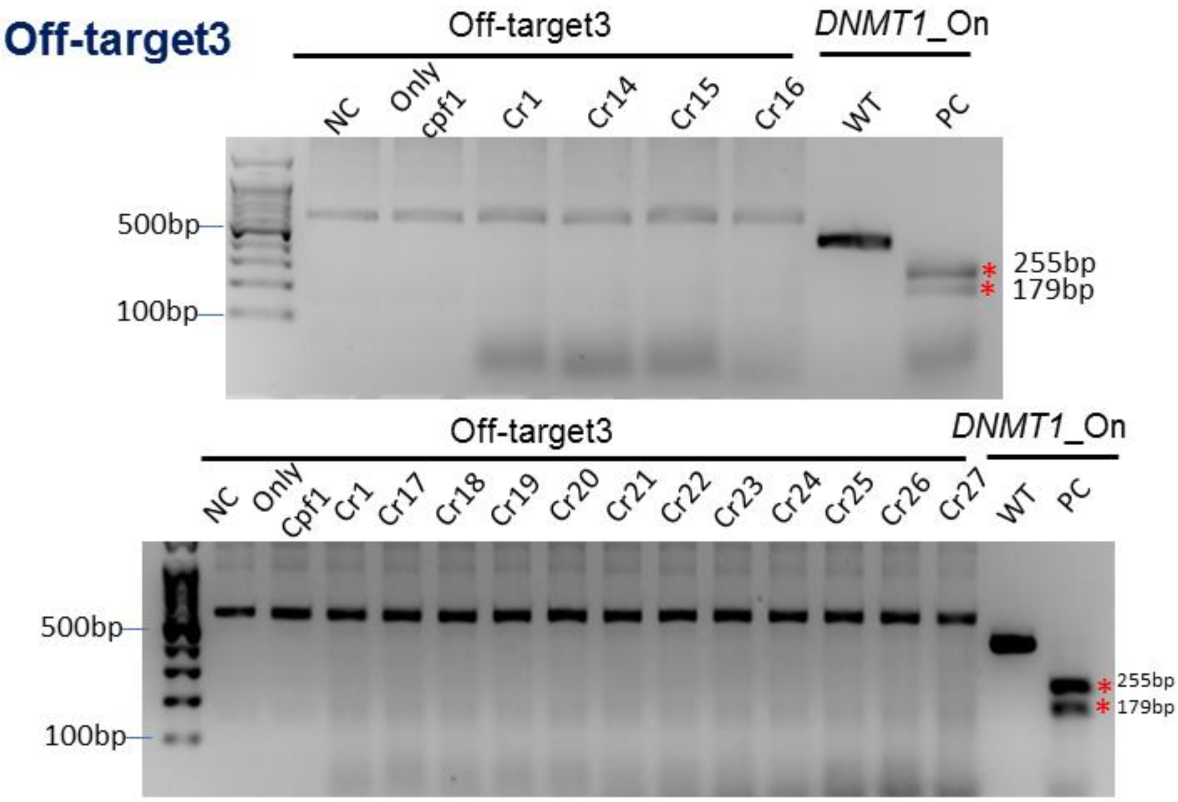
Off-target3
Off-target3
DNMT1_On
NC
Only cpf1
Cr1
Cr14
Cr15
Cr16
WT
PC
500bp
100bp
255bp
179bp
Off-target3
DNMT1_On
NC
Only Cpf1
Cr1
Cr17
Cr18
Cr19
Cr20
Cr21
Cr22
Cr23
Cr24
Cr25
Cr26
Cr27
WT
PC
500bp
100bp
255bp
179bp

*CCR5*

### On-target

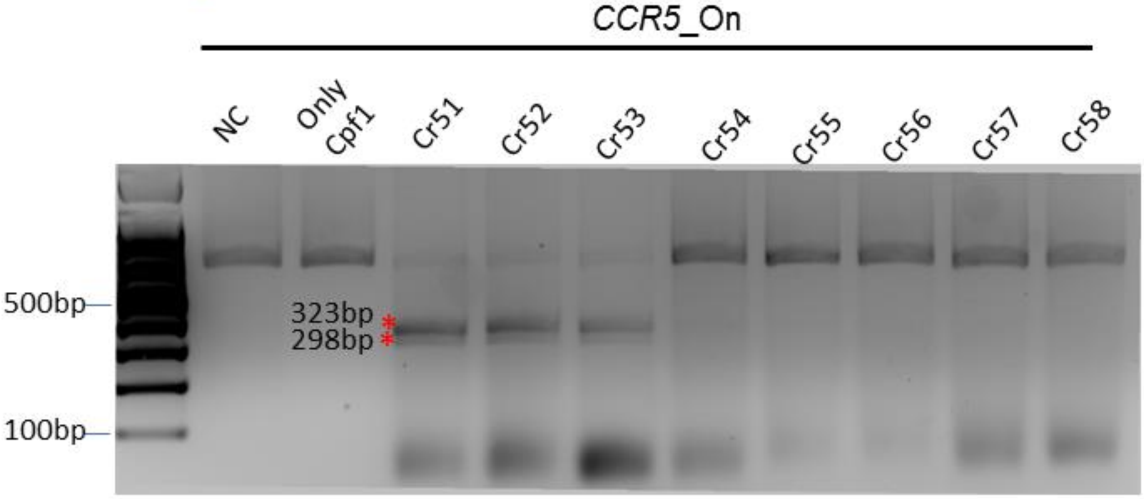


### On-target

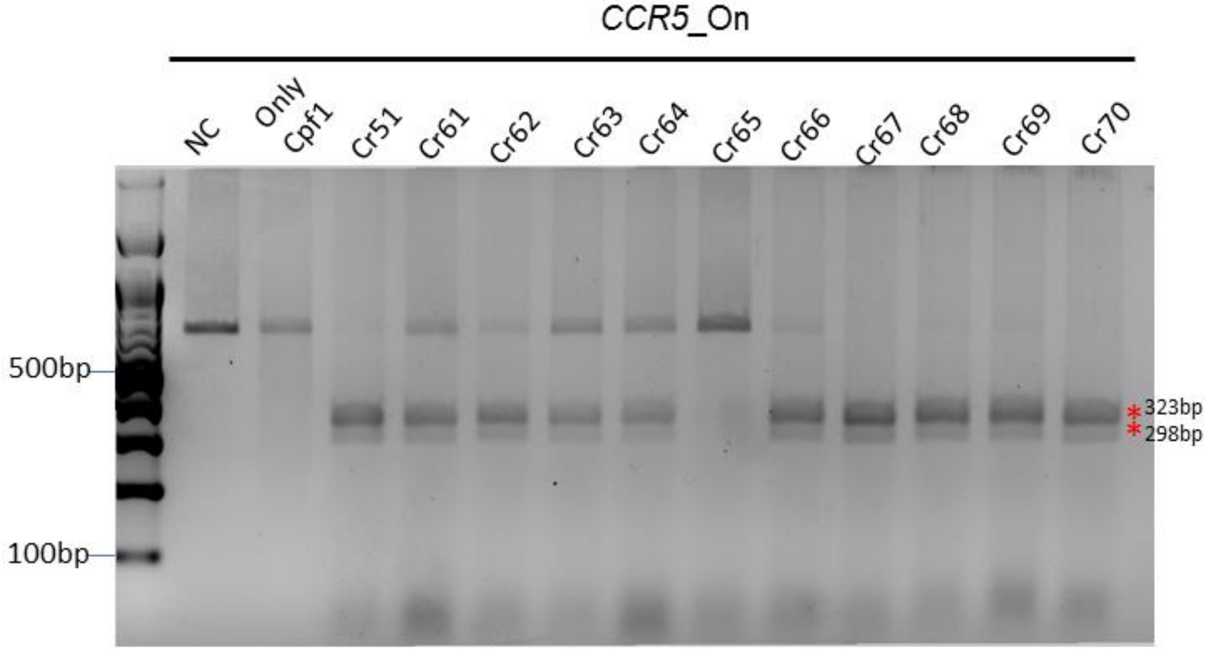

### *FANCF*

#### On-target

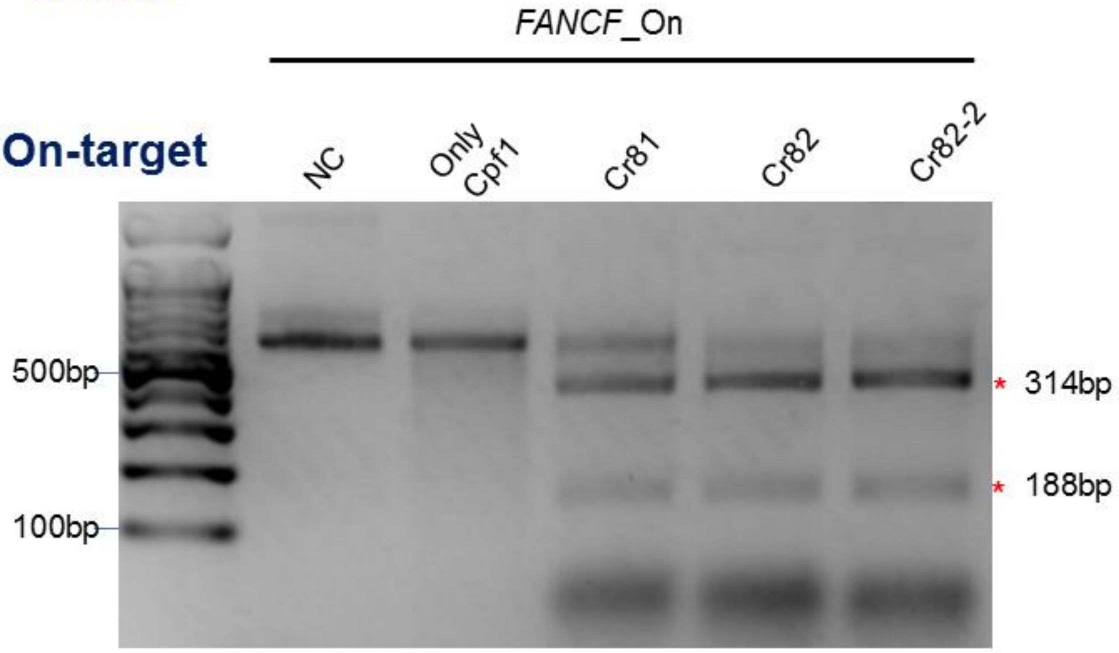


#### Off-target1

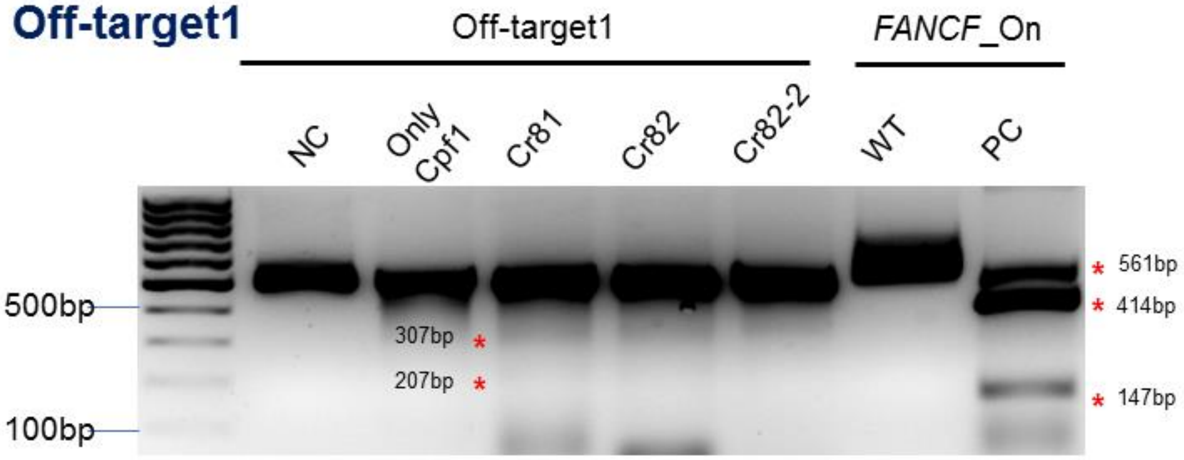


#### Off-target2

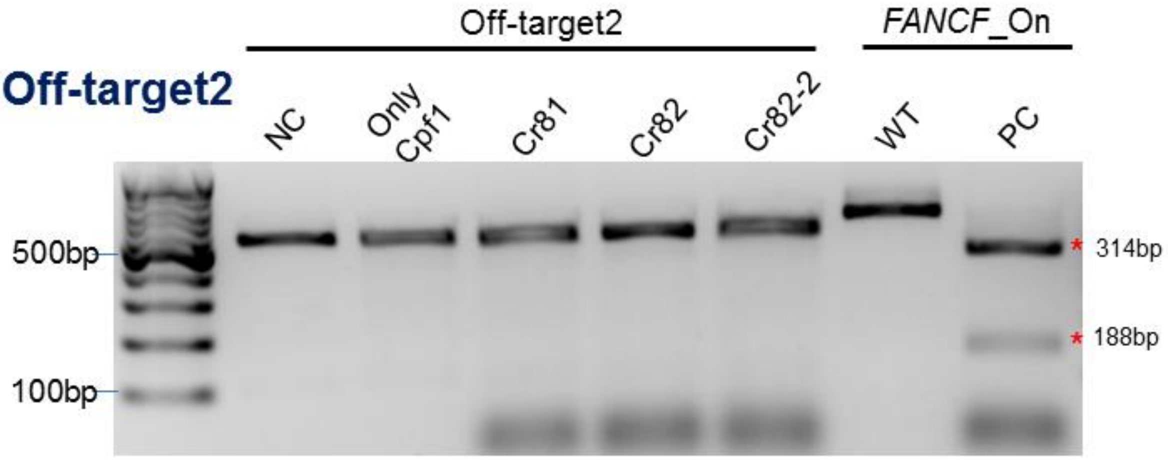

### *GRIN2B*

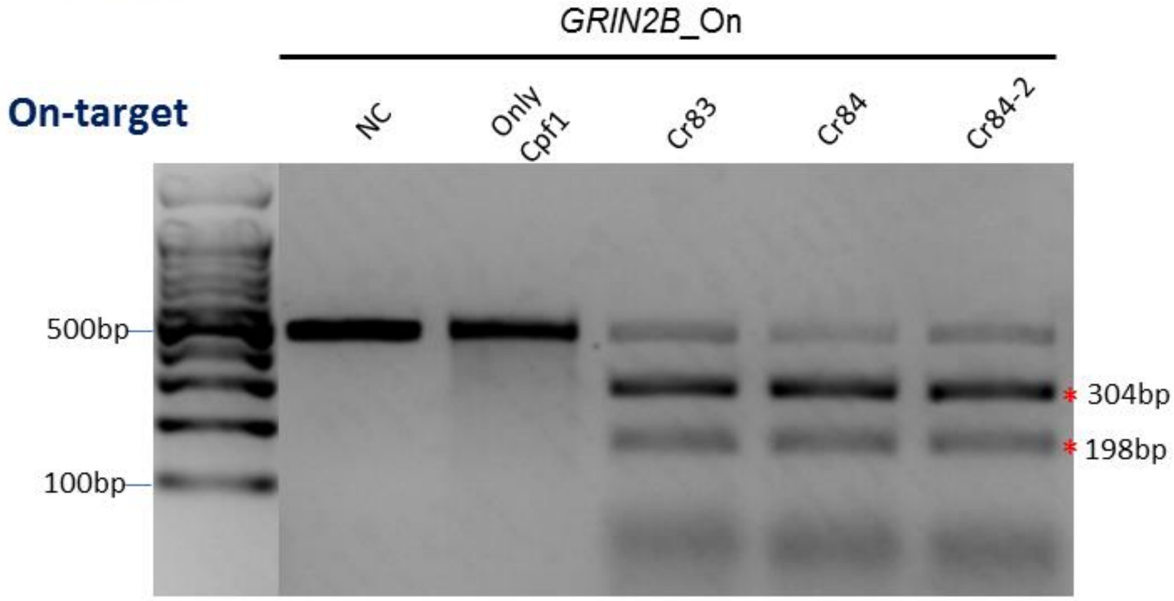
On-target
GRIN2B_On
NC
Only Cpf1
Cr83
Cr84
Cr84-2
500bp
100bp
* 304bp
* 198bp

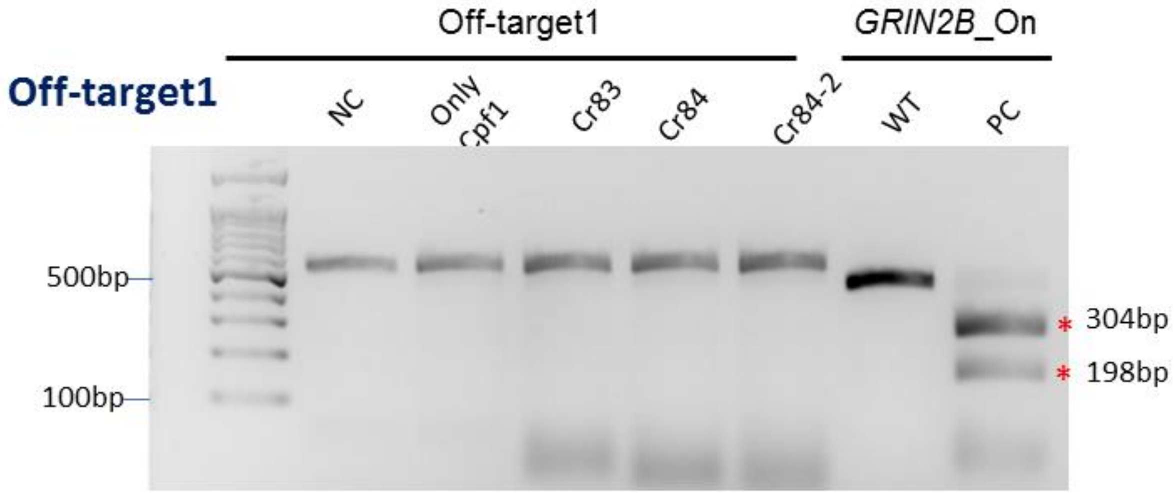
Off-target1
Off-target1
GRIN2B_On
NC
Only Cpf1
Cr83
Cr84
Cr84-2
WT
PC
500bp
100bp
* 304bp
* 198bp

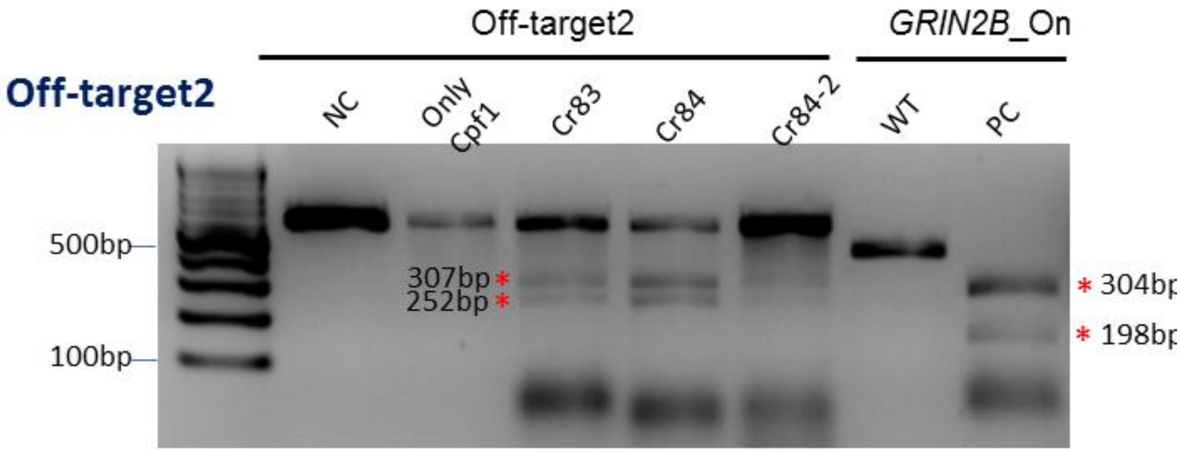
Off-target2
Off-target2
GRIN2B_On
NC
Only Cpf1
Cr83
Cr84
Cr84-2
WT
PC
500bp
100bp
307bp *
252bp *

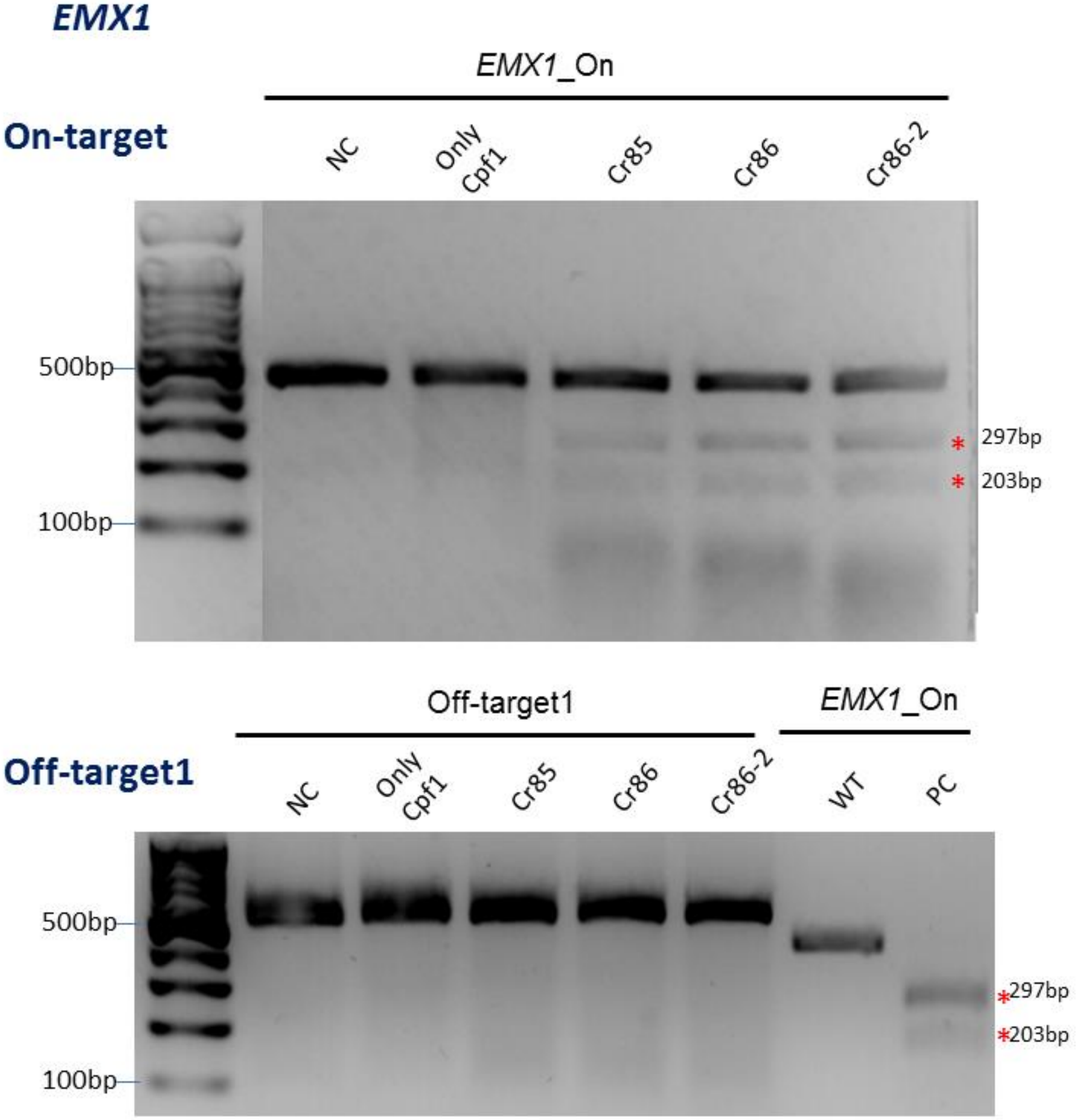


**[Figure S1] Result of the *in-vitro* on/off-target DNA amplicon cleavage assay**

**for various target gene sequence.** PCR amplicons are obtained from purified genomic DNA (HEK293FT) using DNA primers **(Table S2)** corresponding to each target locus (*DNMT1, CCR5, FANCF, GRIN2B, EMX1*). Each amplicon was cleaved and separated on 2% agarose gel. Cleaved DNA fragments are indicated by asterisks.

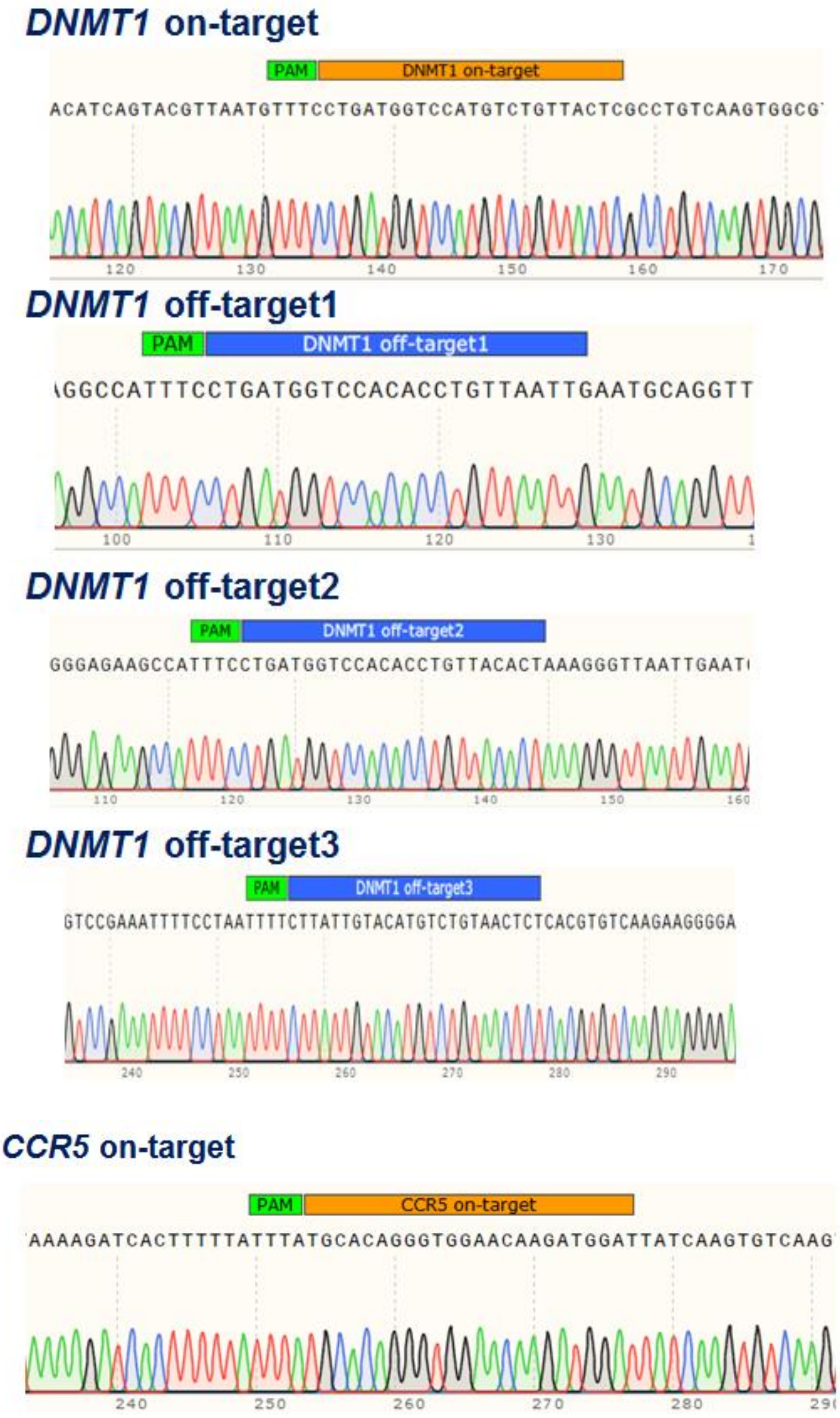

### *FANCF* on-target

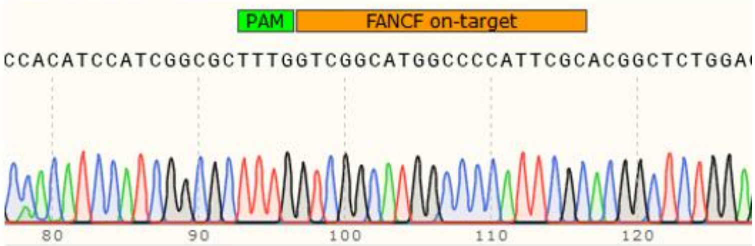


### *FANCF* off-target1

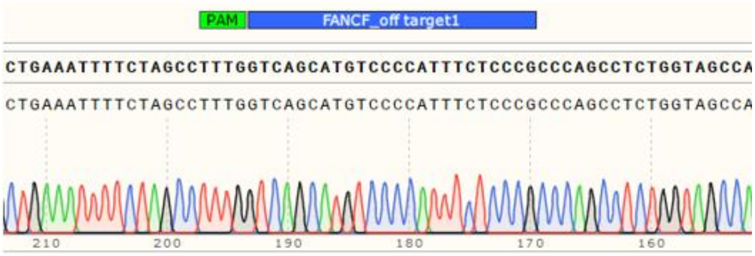


### *FANCF* off-target2

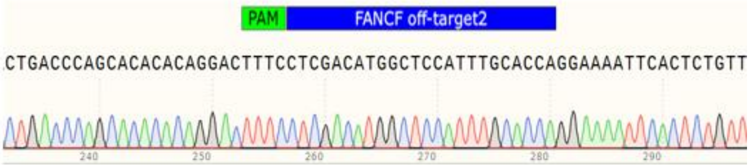


### *GRIN2B* on-target

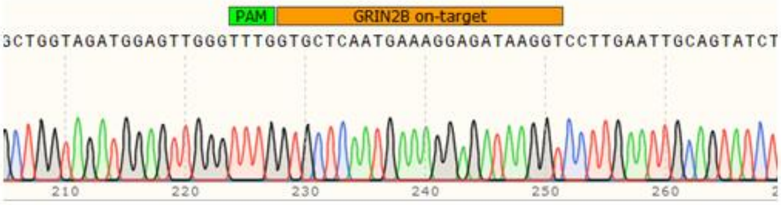


### *GRIN2B* off-target1

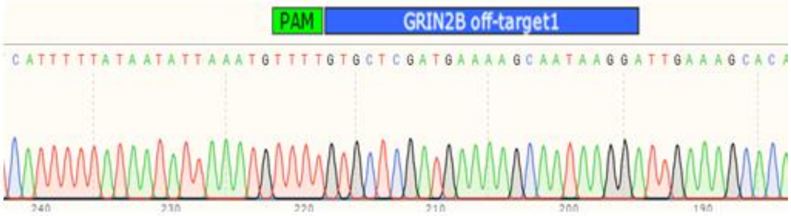


### *GRIN2B* off-target2

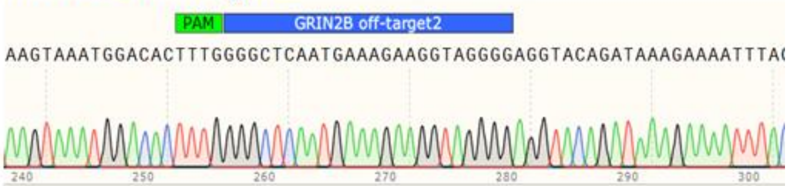

### *EMX1* on-target

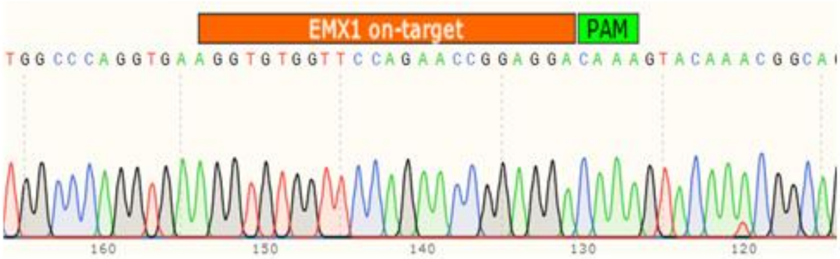


### *EMX1* off-target1

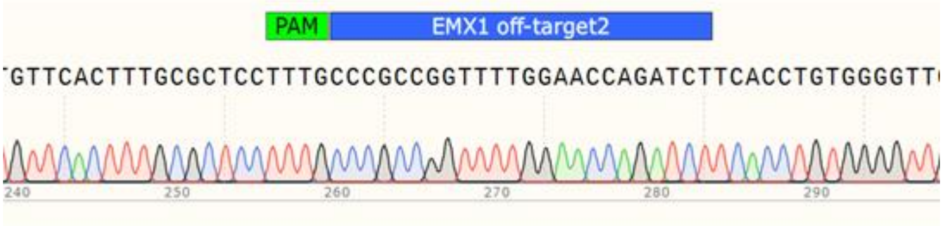


**[Figure S2] Sanger sequencing data from each PCR amplicons of on-target and predicted off-target sites.** PAM sequence (TTTN) for AsCpf1 and on- or off-target sequence in each gene was shown in green, orange and blue color, respectively.

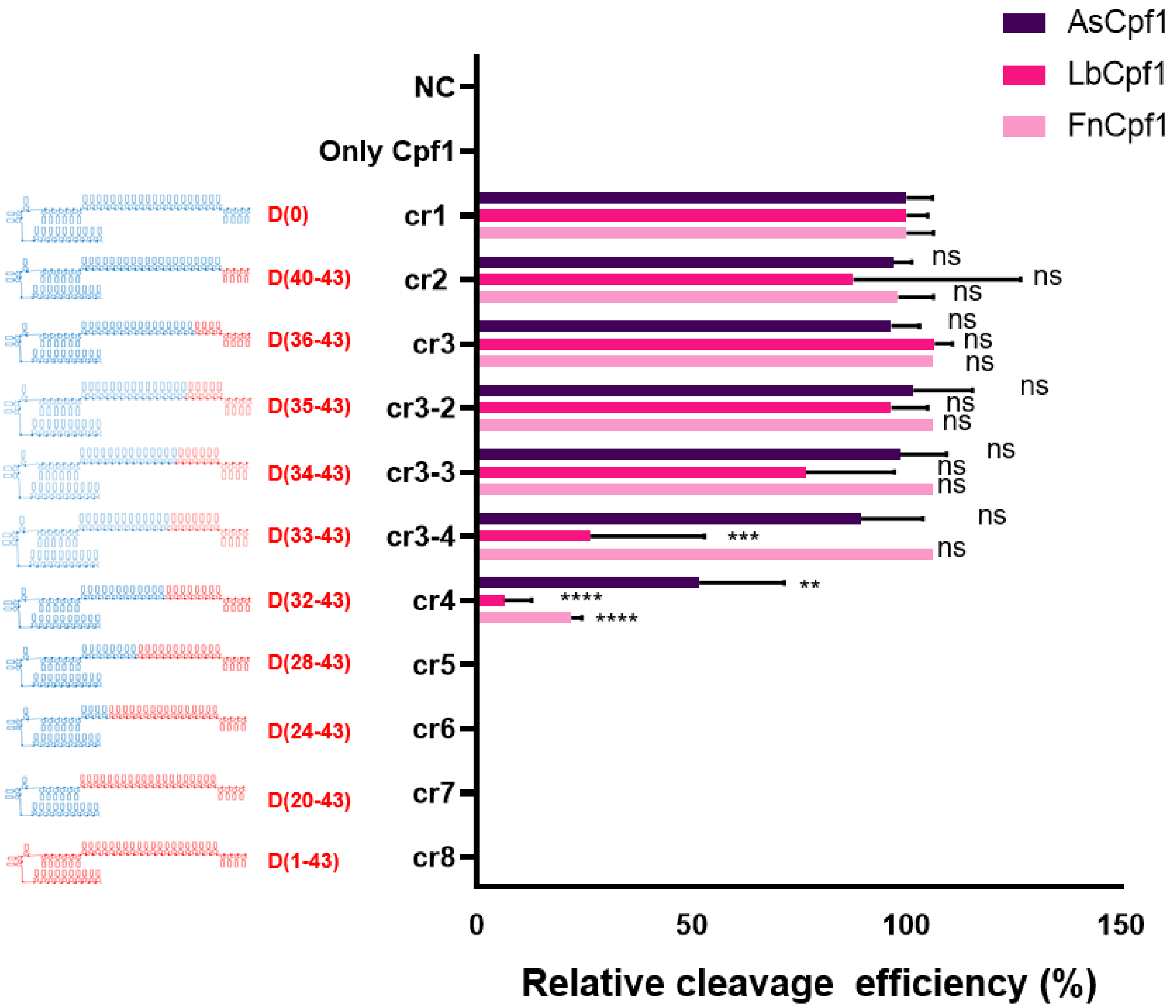


**[Figure S3] Comparison for AsCpf1, LbCpf1 and FnCpf1 cleavage efficiency for *DNMT1* amplicon by using chimeric (cr)RNA.** (cr)RNA of AsCpf1 was partially replaced with DNA from 3'-end. The RNA portion of the (cr)RNA is shown in blue, and the DNA portion is shown in red (number of substituted DNA is indicated) respectively. Cleavage efficiency (%) is a relative ratio among Cpf1 effectors for target DNA cleavage and shown in dark brown, dark pink and pink color. All the cleavage efficiency was calculated from agarose gel separated band intensity (cleavage efficiency (%) = cleaved fragment intensity / total fragment intensity) and normalized to wild-type (cr)RNA (crRNA 1 for *DNMT1*). Data are shown as means ± s.e.m. from three independent experiments. *P*-values are calculated using a one way ANOVA with Tukey's test (ns: not significant, *P**:<0.0332, *P***:<0.0021, *P****:<0.0002, *P*****:<0.0001).

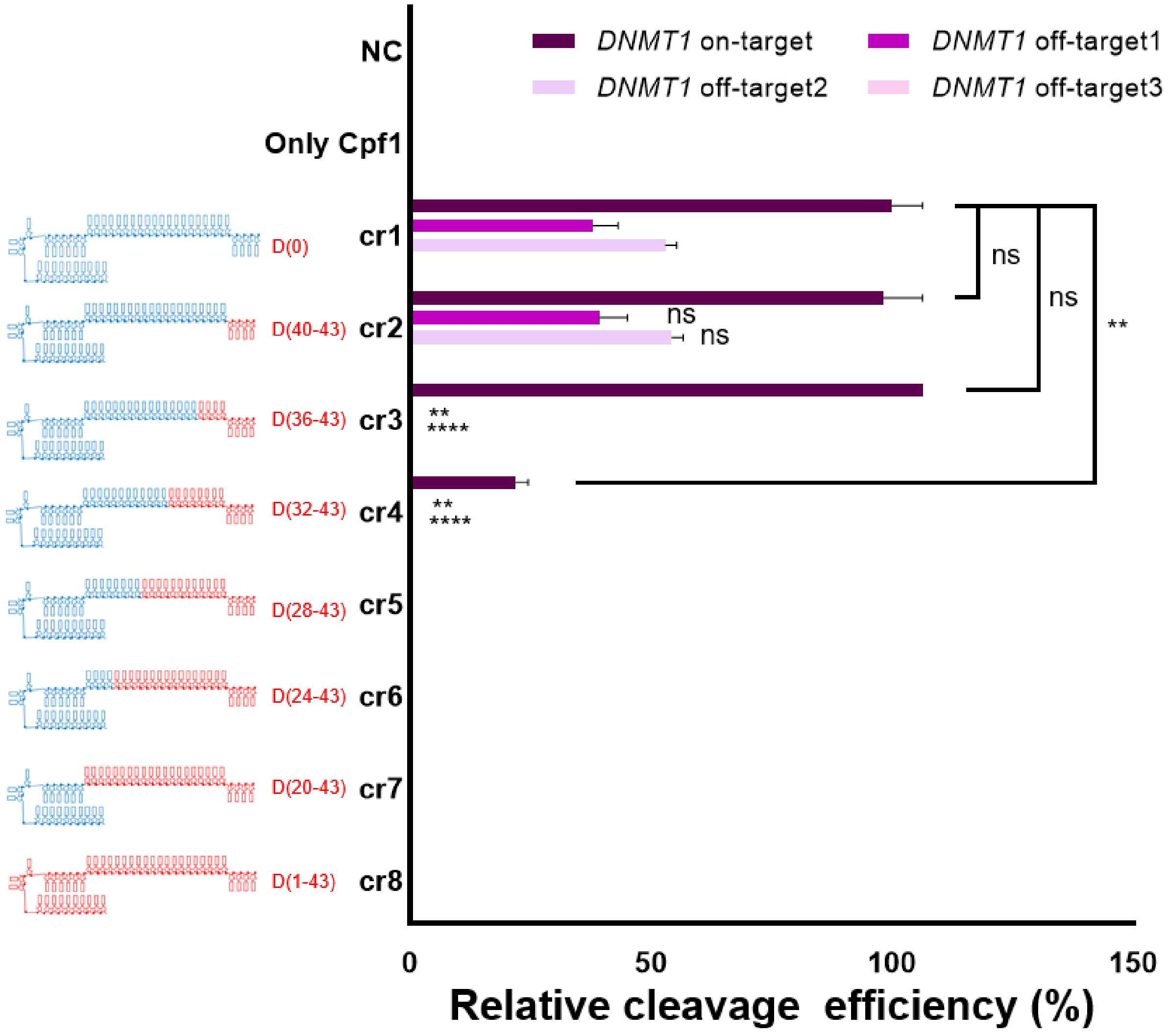


**[Figure S4] Off-target cleavage activity of chimeric (cr)RNA guided FnCpf1.** On/off-target DNA (*DNMT1*: on-target (dark brown), off-target1 (dark purple), off-target2 (light purple), off-target3 (light pink)) amplicon cleavage experiments to confirm the target specificity of FnCpf1 using a chimeric DNA-RNA guide (serial 4nt DNA substitution of (cr)RNA from 3'-end). The RNA portion of the (cr)RNA is shown in blue, and the DNA portion is shown in red (number of substituted DNA is indicated) respectively. Cleavage efficiency is a relative ratio among Cpf1 effectors for target DNA cleavage. All the cleavage efficiency was calculated from agarose gel separated band intensity (cleavage efficiency (%) = cleaved fragment intensity / total fragment intensity) and normalized to wild-type (cr)RNA (crRNA 1 for *DNMT1*). Data are shown as means ± s.e.m. from three independent experiments. *P*-values are calculated using a two-tailed Student's t-test (ns: not significant, P*:<0.05, P**:<0.01, P***:<0.001, P****:<0.0001).

```
CrRNA_1 Indel Pattern
TTTCCTGATGGTCCATGTCTGTTACTCGCCTGTCAAGTGGCGTGACACCG
TTTCCTGATGGTCCATGT------CTCGCCTGTCAAGTGGCGTGACACCG (6bp Del)
TTTCCTGATGGTCCATG-----TACTCGCCTGTCAAGTGGCGTGACACCG (5bp Del)
TTTCCTGATGGTCCA-------TACTCGCCTGTCAAGTGGCGTGACACCG (6bp Del)
TTTCCTGATGGT----------------CCTGTCAAGTGGCGTGACACCG (16bp Del)

CrRNA_2 Indel Pattern
TTTCCTGATGGTCCATGTCTGTTACTCGCCTGTCAAGTGGCGTGACACCG
TTTCCTGATGGTCCATGT------CTCGCCTGTCAAGTGGCGTGACACCG (6bp Del)
TTTCCTGATGGTCCATG-----TACTCGCCTGTCAAGTGGCGTGACACCG (5bp Del)
TTTCCTGATGGTCCATG--------TCGCCTGTCAAGTGGCGTGACACCG (8bp Del)
TTTCCTGATGGT----------------CCTGTCAAGTGGCGTGACACCG (16bp Del)

CrRNA_3 Indel Pattern
TTTCCTGATGGTCCATGTCTGTTACTCGCCTGTCAAGTGGCGTGACACCG
TTTCCTGATGGTCCATGT------CTCGCCTGTCAAGTGGCGTGACACCG (6bp Del)
TTTCCTGATGGTCCATG-------CTCGCCTGTCAAGTGGCGTGACACCG (7bp Del)
TTTCCTGATGGTCCATG--------TCGCCTGTCAAGTGGCGTGACACCG (8bp Del)
TTTCCTGATGGTCCATG-----TACTCGCCTGTCAAGTGGCGTGACACCG (5bp Del)

CrRNA_9 Indel Pattern
TTTCCTGATGGTCCATGTCTGTTACTCGCCTGTCAAGTGGCGTGACACCG
TTTCCTGATGGTCCA----------TCGCCTGTCAAGTGGCGTGACACCG (10bp Del)
TTTCCTGATGGTCCATG-------CTCGCCTGTCAAGTGGCGTGACACCG (7bp Del)
TTTCCTGATGGTCCAT--------CTCGCCTGTCAAGTGGCGTGACACCG (8bp Del)
TTTCCTGATGGTCCATG--------TCGCCTGTCAAGTGGCGTGACACCG (8bp Del)

CrRNA_14 Indel Pattern
TTTCCTGATGGTCCATGTCTGTTACTCGCCTGTCAAGTGGCGTGACACCG
TTTCCTGATGGTCCA-------TACTCGCCTGTCAAGTGGCGTGACACCG (7bp Del)
TTTCCTGATGGTCCATG--------TCGCCTGTCAAGTGGCGTGACACCG (8bp Del)
TTTCCTGATGGTCCATGT---------GCCTGTCAAGTGGCGTGACACCG (9bp Del)
TTTCCTGATGGT----------------CCTGTCAAGTGGCGTGACACCG (16bp Del)

CrRNA_17 Indel Pattern
TTTCCTGATGGTCCATGTCTGTTACTCGCCTGTCAAGTGGCGTGACACCG
TTTCCTGATGGTCCATGT------CTCGCCTGTCAAGTGGCGTGACACCG (6bp Del)
TTTCCTGATGGTCCA-------TACTCGCCTGTCAAGTGGCGTGACACCG (7bp Del)
TTTCCTGATGGTCCAT--------CTCGCCTGTCAAGTGGCGTGACACCG (8bp Del)
TTTCCTGATGGTCCA------TTACTCGCCTGTCAAGTGGCGTGACACCG (6bp Del)

CrRNA_19 Indel Pattern
TTTCCTGATGGTCCATGTCTGTTACTCGCCTGTCAAGTGGCGTGACACCG
TTTCCTGATGGTCCAT--------CTCGCCTGTCAAGTGGCGTGACACCG (8bp Del)
TTTCCTGATGGTCCATG-------CTCGCCTGTCAAGTGGCGTGACACCG (7bp Del)
TTTCCTGATGGTCCATGTCTGTTA--------TCAAGTGGCGTGACACCG (8bp Del)
TTTCCTGATGGTCCATG-----TACTCGCCTGTCAAGTGGCGTGACACCG (5bp Del)

CrRNA_20 Indel Pattern
TTTCCTGATGGTCCATGTCTGTTACTCGCCTGTCAAGTGGCGTGACACCG
TTTCCTGATGGTCCATGT------CTCGCCTGTCAAGTGGCGTGACACCG (6bp Del)
TTTCCTGATGGTCCA------TTACTCGCCTGTCAAGTGGCGTGACACCG (6bp Del)
TTTCCTGATGGTCCAT--------CTCGCCTGTCAAGTGGCGTGACACCG (8bp Del)
TTTCCTGATGGTCCATG--------TCGCCTGTCAAGTGGCGTGACACCG (8bp Del)

CrRNA_22 Indel Pattern
TTTCCTGATGGTCCATGTCTGTTACTCGCCTGTCAAGTGGCGTGACACCG
TTTCCTGATGGTCCA-------TACTCGCCTGTCAAGTGGCGTGACACCG (7bp Del)
TTTCCTGATGGT----------------CCTGTCAAGTGGCGTGACACCG (16bp Del)
TTTCCTGATGGTCCATG--------TCGCCTGTCAAGTGGCGTGACACCG (8bp Del)
TTTCCTGATGGTCCATGT------CTCGCCTGTCAAGTGGCGTGACACCG (6bp Del)

CrRNA_24 Indel Pattern
TTTCCTGATGGTCCATGTCTGTTACTCGCCTGTCAAGTGGCGTGACACCG
TTTCCTGATGGTCCAT--------CTCGCCTGTCAAGTGGCGTGACACCG (8bp Del)
TTTCCTGATGGTCCATGT------CTCGCCTGTCAAGTGGCGTGACACCG (6bp Del)
TTTCCTGATGGTCCA-------TACTCGCCTGTCAAGTGGCGTGACACCG (7bp Del)
TTTCCTGATGGTCCA------TTACTCGCCTGTCAAGTGGCGTGACACCG (6bp Del)

CrRNA_25 Indel Pattern
TTTCCTGATGGTCCATGTCTGTTACTCGCCTGTCAAGTGGCGTGACACCG
TTTCCTGATGGTCCATGT------CTCGCCTGTCAAGTGGCGTGACACCG (6bp Del)
TTTCCTGATGGTCCA-------TACTCGCCTGTCAAGTGGCGTGACACCG (7bp Del)
TTTCCTGATGGTCCATG--------TCGCCTGTCAAGTGGCGTGACACCG (8bp Del)
TTTCCTGATGGTCCATG-----TACTCGCCTGTCAAGTGGCGTGACACCG (5bp Del)

CrRNA_26 Indel Pattern
TTTCCTGATGGTCCATGTCTGTTACTCGCCTGTCAAGTGGCGTGACACCG
TTTCCTGATGGTCCATGT------CTCGCCTGTCAAGTGGCGTGACACCG (6bp Del)
TTTCCTGATGGTCCAT--------CTCGCCTGTCAAGTGGCGTGACACCG (8bp Del)
TTTCCTGATGGTCCA-------TACTCGCCTGTCAAGTGGCGTGACACCG (7bp Del)
TTTCCTGATGGT----------------CCTGTCAAGTGGCGTGACACCG (16bp Del)

CrRNA_27 Indel Pattern
TTTCCTGATGGTCCATGTCTGTTACTCGCCTGTCAAGTGGCGTGACACCG
TTTCCTGATGGTCCAT--------CTCGCCTGTCAAGTGGCGTGACACCG (8bp Del)
TTTCCTGATGGTCCATGT------CTCGCCTGTCAAGTGGCGTGACACCG (6bp Del)
TTTCCTGATGGTCCAT-----------GCCTGTCAAGTGGCGTGACACCG (11bp Del)
TTTCCTGATGGTCCATG--------TCGCCTGTCAAGTGGCGTGACACCG (8bp Del)
```

**[Figure S5] Representative indel patterns from endogenous locus (*DNMT1*) targeted genome editing by chimeric (cr)RNA guided AsCpf1.** Analyzed indel pattern from NGS data of chimeric (cr)RNA guided genome editing in HEK293FT cell. PAM sequence (TTTN) for AsCpf1 is shown in blue and target sequence is shown in red, respectively. Deleted sequence relative to the wild-type reference sequence is indicated by dashed line.

```
CrRNA_1 Indel Pattern
TTTCCTGATGGTCCATGTCTGTTACTCGCCTGTCAAGTGGCGTGACACCG
TTTCCTGATGGTCCATGT------CTCGCCTGTCAAGTGGCGTGACACCG (6bp Del)
TTTCCTGATGGTCCATG-----TACTCGCCTGTCAAGTGGCGTGACACCG (5bp Del)
TTTCCTGATGGTCCA-------TACTCGCCTGTCAAGTGGCGTGACACCG (6bp Del)
TTTCCTGATGGT----------------CCTGTCAAGTGGCGTGACACCG (16bp Del)

CrRNA_2 Indel Pattern
TTTCCTGATGGTCCATGTCTGTTACTCGCCTGTCAAGTGGCGTGACACCG
TTTCCTGATGGTCCATGT------CTCGCCTGTCAAGTGGCGTGACACCG (6bp Del)
TTTCCTGATGGTCCATG-----TACTCGCCTGTCAAGTGGCGTGACACCG (5bp Del)
TTTCCTGATGGTCCATG--------TCGCCTGTCAAGTGGCGTGACACCG (8bp Del)
TTTCCTGATGGT----------------CCTGTCAAGTGGCGTGACACCG (16bp Del)

CrRNA_3 Indel Pattern
TTTCCTGATGGTCCATGTCTGTTACTCGCCTGTCAAGTGGCGTGACACCG
TTTCCTGATGGTCCATGT------CTCGCCTGTCAAGTGGCGTGACACCG (6bp Del)
TTTCCTGATGGTCCATG-------CTCGCCTGTCAAGTGGCGTGACACCG (7bp Del)
TTTCCTGATGGTCCATG--------TCGCCTGTCAAGTGGCGTGACACCG (8bp Del)
TTTCCTGATGGTCCATG-----TACTCGCCTGTCAAGTGGCGTGACACCG (5bp Del)

CrRNA_2-PS Indel Pattern
TTTCCTGATGGTCCATGTCTGTTACTCGCCTGTCAAGTGGCGTGACACCG
TTTCCTGATGGTCCATGT------CTCGCCTGTCAAGTGGCGTGACACCG (6bp Del)
TTTCCTGATGGTCCATG--------TCGCCTGTCAAGTGGCGTGACACCG (8bp Del)
TTTCCTGATGGTCCAT--------CTCGCCTGTCAAGTGGCGTGACACCG (8bp Del)
TTTCCTGATGGTCCA-------TACTCGCCTGTCAAGTGGCGTGACACCG (7bp Del)

CrRNA_3-PS Indel Pattern
TTTCCTGATGGTCCATGTCTGTTACTCGCCTGTCAAGTGGCGTGACACCG
TTTCCTGATGGTCCA---------------TGTCAAGTGGCGTGACACCG (15bp Del)
TTTCCTGATGGTCCATG--------TCGCCTGTCAAGTGGCGTGACACCG (8bp Del)
TTTCCTGATGGTCCA-------TACTCGCCTGTCAAGTGGCGTGACACCG (7bp Del)
TTTCCTGATGGTCCATGTCTGTTAC---CCTGTCAAGTGGCGTGACACCG (3bp Del)
```

**[Figure S6] Representative indel patterns from endogenous locus (*DNMT1*)**

**targeted genome editing by 3'-end modified chimeric (cr)RNA.** Analyzed indel pattern from NGS data of 3'-end chemically modified chimeric (cr)RNA guided genome editing in HEK293FT cell. PAM sequence (TTTN) for AsCpf1 is shown in blue and target sequence is shown in red, respectively. Deleted sequence relative to the wild-type reference sequence is indicated by dashed line.

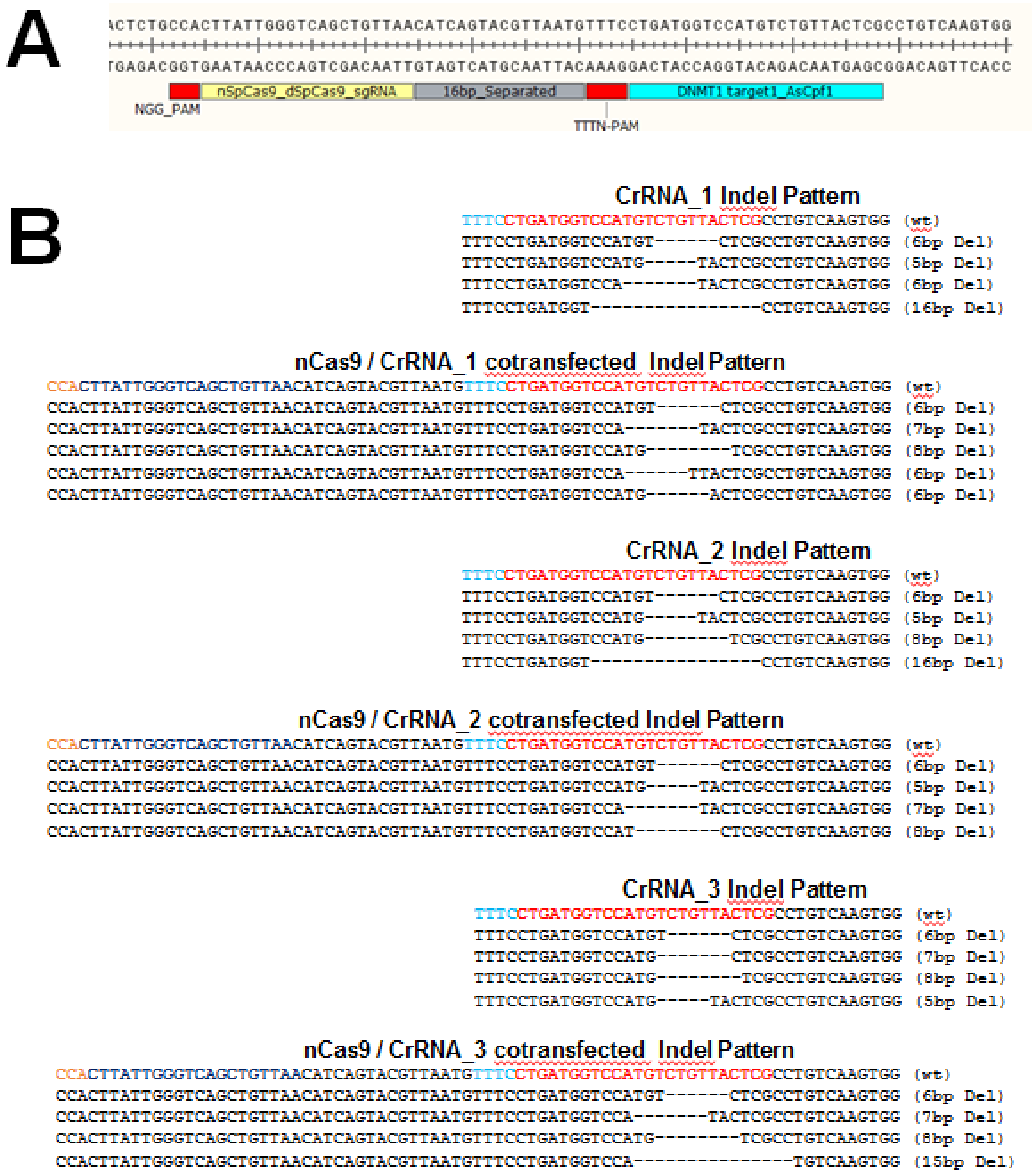


**[Figure S7] *DNMT1* locus targeted genome editing with combination of SpCas9 nickase (D10A) and chimeric (cr)RNA guided AsCpf1 (A)** Targeted endogenous locus of *DNMT1* gene by AsCpf1 and dead(D10A, H840A) or nickase (D10A) type of SpCas9 effectors. PAM sequences (NGG, TTTN) for SpCas9 and Cpf1 effector is shown in red and each target sequence is shown in yellow and cyon color, respectively. **(B)** Representative indel pattern from NGS analysis of chimeric (cr)RNA and nickase guided genome editing in HEK293FT cell. PAM sequence (TTTN) for AsCpf1 is shown in cyon and target sequence is shown in red. PAM sequence (NGG) for SpCas9 is shown in orange and target sequence is shown in dark blue, respectively. Deleted sequence relative to the wild-type reference sequence is indicated by dashed line.

A

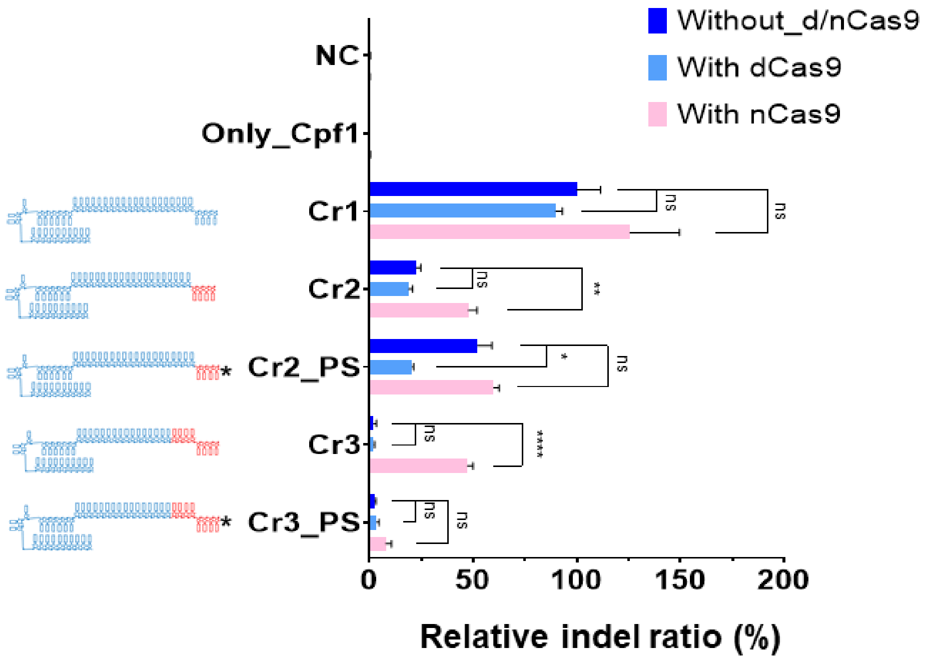


B

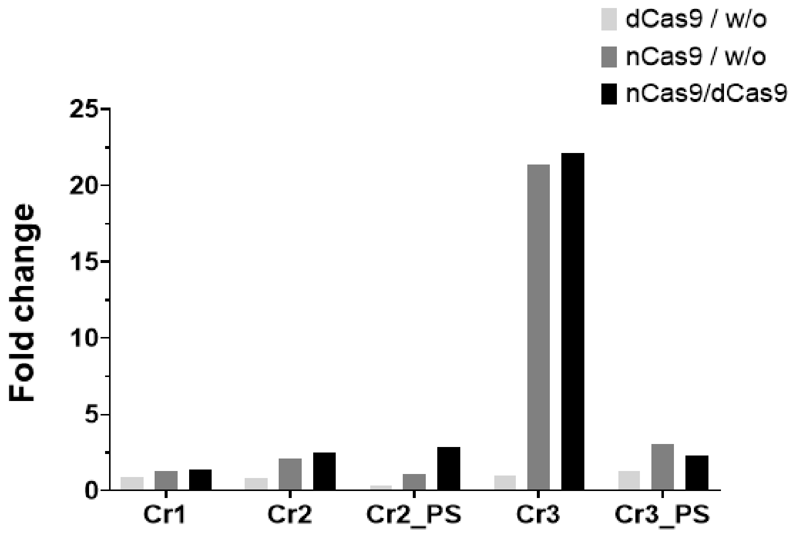


**[Figure S8] Endogenous locus (*DNMT1*) targeted genome editing with combination of 3’-Phosphorothioate (PS) modified chimeric (cr)RNA and**

**SpCas9 nickase. (A)** Intracellular *DNMT1* gene editing in HEK293FT cell using 3′-end PS-modified chimeric DNA-RNA guided Cpf1 and SpCas9 (D10A) nickase combination. X axis indicates the relative indel frequency (%). Only (cr)RNA-Cpf1 treated (dark blue) and combination with dCas9 (blue) or nCas9 (pink) treated samples were indicated. Data are shown as means ± s.e.m. from three independent experiments. *P*-values are calculated using a Student's t-test (ns: not significant, P*:<0.05, P**:<0.01, P***:<0.001, P****:<0.0001). **(B)** Fold increase of the relative indel ratios (%) among only (cr)RNA-Cpf1 treatment and additional dCas9 or nCas9 treated samples were indicated. dCas9 and nCas9 indicates deactivated Cas9 (D10A, H840A) and nickase Cas9 (D10A), respectively.

## A

| *DNMT1* target | Mismatch No. | Sequence |
|---|---|---|
| On-target | 0 | TTTCCTGATGGTCCATGTCTGTTACTCG |
| Off-target1 | 5 | TTTCCTGATGGTCCACACCTGTTAATTG |
| Off-target2 | 5 | TTTCCTGATGGTCCACACCTGTTACACT |
| Off-target3 | 6 | TTTTCTTATTGTACATGTCTGTAACTCT |

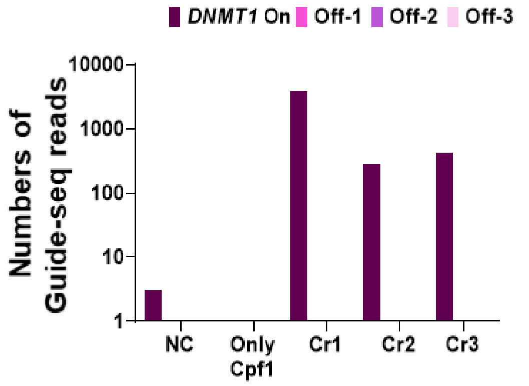


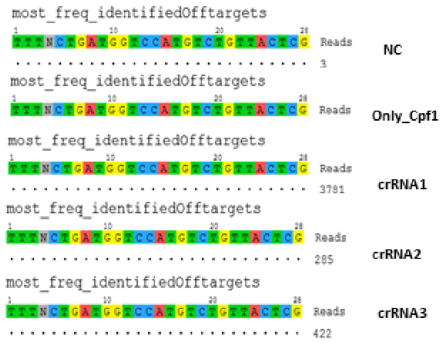


## B

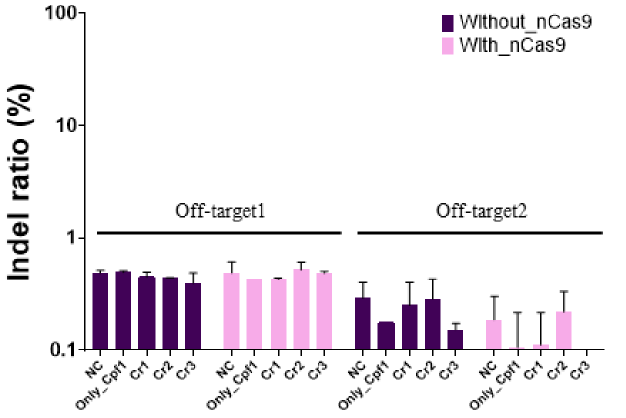


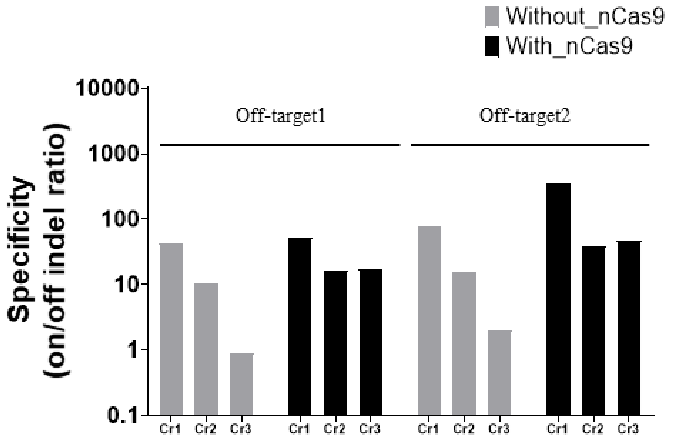

**[Figure S9] Off-target mutation analysis for *DNMT1* site targeted with combination of SpCas9 nickase(D10A) and chimeric (cr)RNA guided Cpf1. (A)** Top: Table shows selected off-target site from Guide-seq analysis. On-and off-target sequence information for *DNMT1* gene is shown. The PAM sequence is underlined and shown in bold. Mismatched nucleotides to reference sequence are shown in red. Bottom: Genome-wide off-target analysis with Guide-seq for *DNMT1* target site in **(Figure 7A)**. Numbers of Guide-seq reads were indicated with different colors (dark brown: on-target, dark pink: off-target 1, purple: off-target 2, light pink: off-target 3). **(B)** Off-target indel ratio (%) for site1 and site2 was calculated from targeted amplicon sequencing data in **(Figure 7A)**. Data are shown as means ± s.e.m. from three independent experiments. *P*-values are calculated using a Student's t-test (ns: not significant, P*:<0.05, P**:<0.01, P***:<0.001, P****:<0.0001).

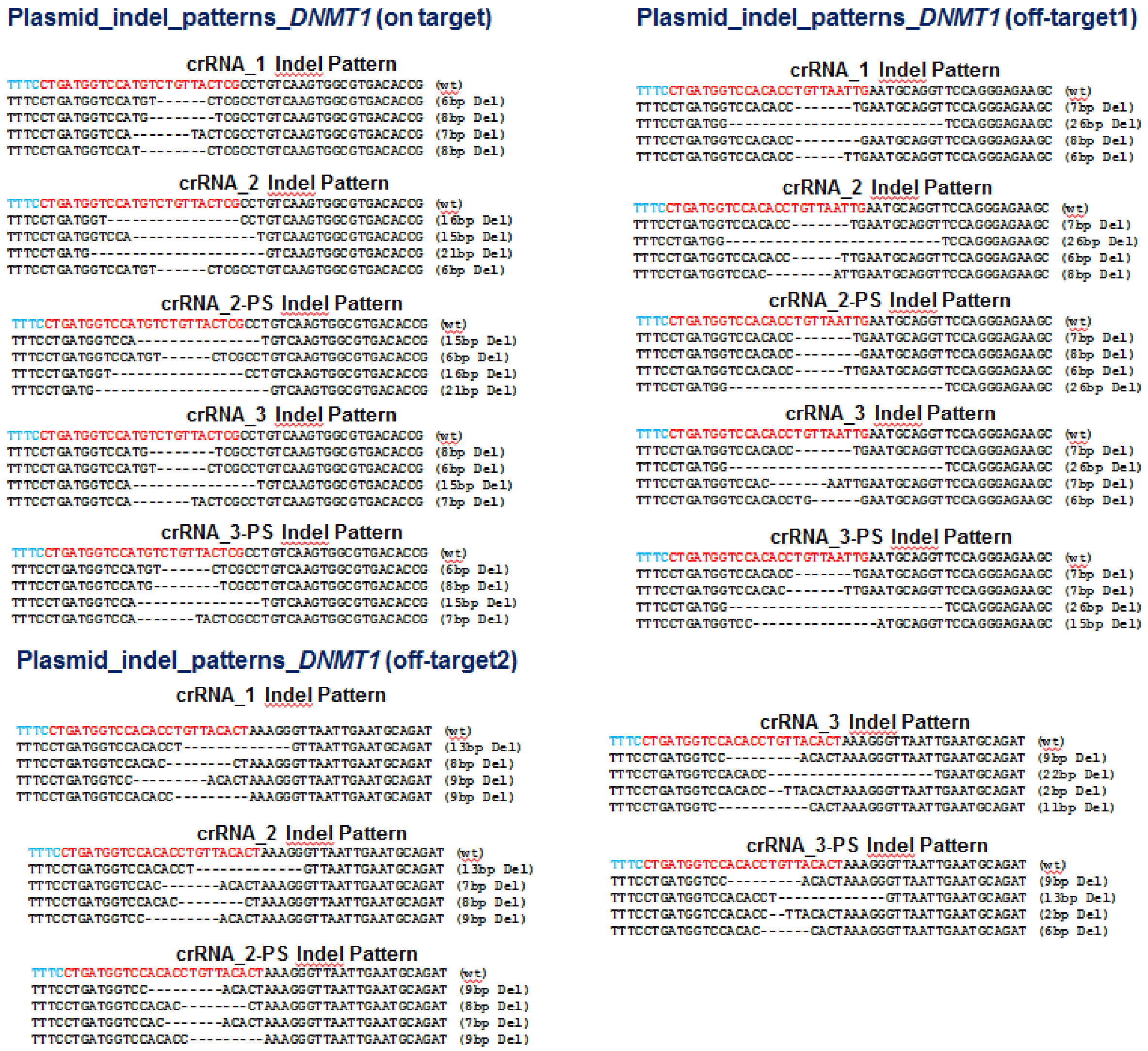


**[Figure S10] Representative indel patterns from plasmid (*DNMT1* on-, off-target1, off-target2) targeted genome editing by chimeric (cr)RNA guided AsCpf1.** Analyzed indel pattern from NGS data of 3'-end chemically modified chimeric (cr)RNA guided genome editing in HEK293FT cell. PAM sequence (TTTN) for AsCpf1 is shown in blue and target sequence is shown in red, respectively. Deleted sequence relative to the wild-type reference sequence is indicated by dashed line.
